# *Mycobacterium tuberculosis* Ku stimulates multi-round DNA unwinding by UvrD1 monomers

**DOI:** 10.1101/2023.09.29.560180

**Authors:** Ankita Chadda, Alexander G. Kozlov, Binh Nguyen, Timothy M. Lohman, Eric A. Galburt

## Abstract

*Mycobacterium tuberculosis* is the causative agent of Tuberculosis. During the host response to infection, the bacterium is exposed to both reactive oxygen species and nitrogen intermediates that can cause DNA damage. It is becoming clear that the DNA damage response in *Mtb* and related actinobacteria function via distinct pathways as compared to well-studied model bacteria. For example, we have previously shown that the DNA repair helicase UvrD1 is activated for processive unwinding via redox-dependent dimerization. In addition, mycobacteria contain a homo-dimeric Ku protein, homologous to the eukaryotic Ku70/Ku80 dimer, that plays roles in double-stranded break repair via non-homologous end-joining. Ku has been shown to stimulate the helicase activity of UvrD1, but the molecular mechanism, as well as which redox form of UvrD1 is activated, is unknown. We show here that Ku specifically stimulates multi-round unwinding by UvrD1 monomers which are able to slowly unwind DNA, but at rates 100-fold slower than the dimer. We also demonstrate that the UvrD1 C-terminal Tudor domain is required for the formation of a Ku-UvrD1 protein complex and activation. We show that *Mtb* Ku dimers bind with high nearest neighbor cooperativity to duplex DNA and that UvrD1 activation is observed when the DNA substrate is bound with two or three Ku dimers. Our observations reveal aspects of the interactions between DNA, *Mtb* Ku, and UvrD1 and highlight the potential role of UvrD1 in multiple DNA repair pathways through different mechanisms of activation.

**Declaration of Interests**

None

## INTRODUCTION

About one-third of the human population is estimated to be infected with *Mycobacterium tuberculosis* (*Mtb*), the causative agent of Tuberculosis [1]. During infection, the host immune response exposes *Mtb* to reactive oxygen species and reactive nitrogen intermediates that can lead to DNA damage [2,3]. This damage occurs in the form of base modifications, single-strand breaks, and double-strand breaks (DSB) [4,5]. While single-strand breaks can be bypassed by replication and transcription machinery, specialized enzymes are required to repair DSBs [6,7]. Mycobacteria has three known DSB repair pathways: single-strand annealing (SSA), homologous recombination (HR), and nonhomologous end joining (NHEJ) [8,9]. Repair can proceed through SSA if repeating homologous sequences are available on each side of the break. Resection leads to complementary single strands that can anneal and subsequent flap removal and end-joining lead to repair with deletion of the resected sequences [8]. HR is a high-fidelity DNA repair pathway but can only function when a second intact DNA copy is present; for example, during or after replication and before division [10,11]. NHEJ can either be error-free (if the ends are sealed directly) or mutagenic depending on how the DNA ends are processed prior to ligation [9]. Since NHEJ acts as the major pathway in non-replicating cells, it may be of significant relevance in persistence and pathogenesis of *Mtb* [12–14].

Prokaryotic NHEJ is accomplished through the action of two proteins: Ku and Ligase D (LigD). Bacterial Ku proteins are ∼30 kDa, form homodimers, bind double-stranded DNA, and are responsible for identifying, binding, and protecting DSBs [15,16]. The core domain of bacterial Ku is homologous to that of the eukaryotic Ku70:Ku80 heterodimer and is highly conserved among bacterial Ku’s from different species. Each core domain forms one-half of a dimeric ring-like structure that encircles DNA [17–19]. In contrast, the C-terminal domain (CTD) is unique to bacterial Ku and consists of 20-25 conserved amino acids and an extended region of basic residues of varying lengths (i.e., 14 in *Mtb* to 40 in *M. smegmatis*). The conserved sequence is important for recruiting LigD to DNA ends and the extended basic region has also been implicated in DNA binding and threading [20].

DNA helicases play varied roles in DNA repair including unwinding duplex DNA, removing damaged ssDNA, stimulating the resection of blunt-ended substrates, and removing DNA-bound proteins [21–24]. Many bacterial helicases like UvrD, UvrD1, Rep, and PcrA, and eukaryotic helicases and helicase complexes like TFIIH, FANCJ, and WRN are involved in DNA repair pathways [25–30]. *Mtb* UvrD1 belongs to the UvrD/PcrA subgroup of SF1 super-family helicases and consists of two RecA-like domains (1A and 2A) with two accessory domains (1B and 2B). Members of this subgroup have been shown to unwind DNA as dimers; monomers are ssDNA translocases, but can become helicases through interactions with accessory factors [31–39]. In addition, they sometimes contain a C-terminal Tudor domain which has been implicated in protein-protein interactions with binding partners [40,41]. We previously showed that *Mtb* UvrD1 exists as a mixture of monomers and dimers depending on redox potential [42]. Although both monomers and dimers of UvrD1 can bind and translocate on ssDNA, only the dimer formed in oxidative conditions via a specific disulfide bond between the 2B domains possesses processive helicase activity [42].

Studies performed prior to the elucidation of the redox-dependent dimerization and activation of UvrD1 showed that *Mtb* Ku stimulates the helicase activity of UvrD1 and provided evidence for protein-protein interactions between Ku and the CTD of UvrD1 [43,44]. Here we show that Ku specifically stimulates the helicase activity of UvrD1 monomers, but only under multi-round binding conditions. This activity is slow (∼100-fold slower compared to the dimer) and inefficient (∼15% fraction of DNA substrates are unwound in the presence of saturating Ku). Furthermore, removing the C-terminal Tudor domain of UvrD1 abrogates the Ku-dependent activation. Lastly, *Mtb* Ku coats the dsDNA region of our unwinding substrates even in the presence of 3’ single-stranded overhangs. Ku binding is cooperative and dependent on magnesium and Ku-stimulated UvrD1 unwinding occurs even when multiple Ku dimers are bound to the DNA. These data serve to define the molecular mechanism for Ku-based activation of UvrD1 while quantitatively comparing the slow Ku-stimulated activity of monomeric UvrD1 in multi-round conditions to the rapid and efficient single-round (*i*.*e*., processive) activity of oxidatively formed dimers. These results highlight a possible role of UvrD1 monomers in remodeling and processing Ku-bound DNA during DSB repair.

## MATERIAL AND METHODS

### Protein purification

The open reading frame encoding *M. tuberculosis* Ku (Rv0937c) was cloned in pET45b vector with ampicillin resistance using a BamH1 site at the start codon and a HindIII site 3 ′ of the stop codon. The plasmids were confirmed by sequencing to exclude the acquisition of unwanted coding changes. The pET-*Mtb* Ku plasmid was used to transform *E. coli* BL21(DE3). Ku was overexpressed by induction with 0.5 mM isopropyl-β-D-thiogalactopyranoside (IPTG) at OD of 0.5, followed by incubation at 23°C for 16 h with constant shaking. The cells were harvested by centrifugation, and the pellets were either stored at −80 °C or used for subsequent procedures that were performed at 4 °C. An ∼12 g cell pellet obtained from 1 L of liquid culture was resuspended in lysis buffer (50 mM Tris-HCl, pH 7.5, 0.25 M NaCl, 10% sucrose). The lysates were sonicated, and insoluble material was removed by centrifugation at 14K for 45 minutes. The soluble extracts were applied to 2 ml columns of nickel-nitrilotriacetic acid agarose (Ni-NTA) (QIAGEN catalogue no. 30210) that had been equilibrated with lysis buffer. The columns were washed with 10X column volume of wash buffer (50 mM Tris-HCl, pH 8.0, 0.25 M NaCl,10% glycerol) and then eluted stepwise with wash buffer containing 50-, 100-, 200-, 500-, and 1000-mM imidazole. The polypeptide compositions of the column fractions were monitored by SDS-PAGE. Ku was recovered in 100-and 200 mM imidazole fractions. After overnight dialysis in (50 mM Tris-HCl pH 8.0, 60 mM NaCl, 10% glycerol) the protein was passed through DEAE-Sephacryl chromatography followed by size exclusion chromatography where it eluted as a dimer. *Mtb* UvrD1 proteins both wild type and C-terminal deletion mutant were expressed and purified as described previously [42].

### DNA substrates

Single-stranded DNA, which is either labelled with Cy5, Cy3, FAM or BHQ2 were obtained from Integrated DNA technologies (IDT, Coralville, IA). Duplex DNA substrates were prepared by annealing ssDNA oligos with the fluorescent label at 5’end of the single strand that was mixed with an equimolar concentration of unlabelled, BHQ-2 or FAM labelled complementary strand in 10 mM Tris pH 8.0, 50 mM NaCl, followed by heating to 95 °C for five minutes and slow cooling to room temperature.

### Stopped-flow DNA unwinding assays

All stopped-flow unwinding experiments were carried out at 25°C using an Applied Photophysics instrument SX-20, minimum total shot volume 100 μl, dead time 2 ms. Experiments were carried out in Buffer with Tris pH 8.0, 75 mM NaCl, 5mM MgCl_2_, 20% glycerol and with 1 mM DTT for reducing conditions. The DNA substrates used in the assay are double-stranded 18-basepair or 32-basepair DNA with a dT_20_, dT_40_, or dT_60_ tail with a Cy5 fluorophore attached to the 5’ end of the long strand and a black hole quencher (BHQ-2) attached to the 3’ end of the short strand. A Cy5 fluorophore was excited using a 625 nm LED (Applied Photophysics Ltd., Leatherhead, UK) and fluorescence emission was monitored at wavelengths >665 nm using a long-pass filter (Newport Optics). DNA unwinding was monitored as the increase in Cy5 fluorescence upon DNA strand separation. The traces represent the average of 5 independent shots and at least two different protein preparations. UvrD1 (400 nM) was incubated with or without Ku (1600 nM) and 20 nM DNA in one syringe and rapidly mixed with the contents of the other syringe: containing 2 mM ATP, 10 mM Mg^+2^, and either no TRAP for multi-round experiments or one of three protein TRAPs (an 18 nt ssDNA protein, an 18bp-dT20 partial duplex, or a dT_40_-10bp hairpin) in excess of protein (25X, 5 µM) for single-round experiments. The ssDNA also serves as a trap for any unwound DNA strands. In the case of Ku-stimulated unwinding, the slow rate of DNA unwinding was monitored over 2400 seconds. The unwinding signal was normalized to the signal from positive and negative controls to calculate the fraction of DNA unwound. The signal for fully unwound DNA is alternatively obtained using 20 or 50 nM of the labeled partial duplex denatured in the presence of TRAP or unannealed 20 or 50 nM single-stranded Cy5 labeled DNA in the presence of UvrD1 and Ku. The signal for fully duplex DNA was obtained from 20 nM duplex DNA with Cy5 and the BHQ-2 in the presence of UvrD1 and Ku but without ATP.

### Analysis of Ku binding to the DNA unwinding substrate

Ku protein binding to the DNA unwinding substrate was examined by monitoring the quenching of fluorescence intensity of a 32 bp double-stranded DNA with a dT_20_ single-stranded tail, containing a fluorescein (FAM) fluorophore at the 5’ end of the duplex DNA end. Fluorescence titrations were performed using a PTI QM-2000 spectrofluorometer (Photon Technologies, Inc., Lawrenceville, NJ) at 25.0 °C in buffer containing Tris pH 8.0, 75 mM NaCl, 20% glycerol, 5 mM MgCl_2_, 2 mM DTT and 9 µM BSA. Four titrations were performed in duplicate or triplicate at 10 nM, 30 nM, 100 nM and 300 nM FAM-DNA in a 3 mL quartz cuvette by adding aliquots of Ku protein while monitoring fluorescein fluorescence intensity (excitation and emission wavelengths of 490 and 520 nm, respectively). Corrections for dilution were applied as described [45].

Using these titrations to obtain binding stoichiometries and binding affinities, requires knowledge of the relationship between the observed flurescence quenching and the average extent of Ku binding to the DNA. To accomplish this, plots of fluorescence quenching vs. total Ku dimer concentration at the four DNA contrations were analyzed using the macromolecule binding density method as described [46]to determine the relationship between the observed fluorescence quenching and the average extent of Ku binding to the DNA. Briefly, at any constant value of fluorescence quenching (along any horizontal line intersecting each of the titration curves obtained at the four total DNA concentration ([D]_TOT_), the free Ku dimer concentration must be constant, and thus the average extent of binding (<Ku>=[Ku]_bound_/([D]_TOT_) must also be constant for each of the titrations at the four total DNA concentrations. This yields a set of four pairs of values of [D]_TOT_ and [Ku]_TOT_ for that one horizontal line. Since [Ku]_TOT_ = [Ku]_free_ + <Ku>([D]_TOT_), a plot of [Ku]_TOT_ vs. [D]_TOT_ will be linear, with slope equal to <Ku> and intercept equal to [Ku]_free_ for that horizontal line at that particular value of fluorescence quenching. Repeating this for a series of horizontal lines drawn at different values of fluorescence quenching yields a model free binding isotherm (<Ku> as a function of [Ku]_free_). From this one can also obtain the relationship between the fluorescence quenching signal and the average extent of Ku binding (<Ku>). We found that, within our uncertainties, the average fluorescence quenching was linearly related to <Ku>, hence <Ku> = Q_obs_/Q_max_.

We used a finite lattice nearest neighbour cooperativity model to obtain estimates of the equilibrium binding association constant (K) and nearest neighbour cooperativity (ω) for Ku binding to the DNA substrate for non-specific binding of a large ligand (protein), with occluded site size, n, to a DNA of length L base pairs (bp) [47].

The cooperativity parameter, ω, represents the additional affinity obtained when two Ku dimers bind at contiguous sites on the DNA. From the stoichiometric titration curve performed at 300 nM DNA, we find that 3 Ku dimers can bind to each 32 bp DNA substrate at saturating Ku concentrations. Hence, we assign an occluded site size of n=11 bp. As discussed below, this is only possible if the Ku dimers bind with high positive cooperativity to the DNA lattice. Non-cooperative binding of Ku would result in strong binding of only two Ku dimers to the DNA lattice at the Ku concentrations used in our experiments. To simplify the analysis, we use a DNA length of 33 bp in our modelling, rather than the actual length of 32. Since Ku does not bind to the single stranded DNA flanking the duplex, the first Ku has (L-n+1) = 23 binding sites on the duplex, assuming that Ku must maintain an 11 nucleotide site (*i*.*e*., Ku does not bind to a partial site overhanging the DNA end). Two Ku dimers can bind the DNA in 78 different configurations, 12 with nearest neighbour cooperativity, ω, and 66 non-cooperatively.

The binding polynomial, P, for this system, for L=33 and n=11, with the free DNA as the reference state, is given in Eq. (1),

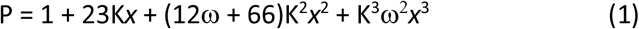

where K is the equilibrium association constant for one Ku dimer binding to the DNA and *x* is the free Ku dimer concentration. The average extent of Ku dimer binding per DNA is obtained as <Ku> = (dlnP/dlnx) and is given in Eq. (2).

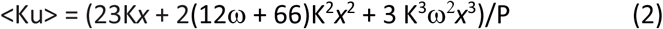

Since we find that <Ku> = Q_obs_/Q_max_, a non-linear least squares fitting of the four titration curves, to Eq. (2) yields estimates of K and ω. The fraction of DNA molecules with one, two or three Ku dimers bound, f_1_, f_2_ and f_3_, respectiviely, can be calculated from Eq. (3-5).

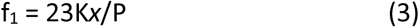

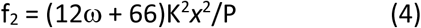

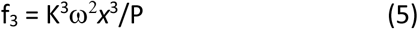

Based on the linear dependence of Q on <Ku>, we assume Q=Q_max_(1/3f_1_+2/3f_2_+f_3_). Hence the final model has only three parameters, K, ω and Q_max_.

### Analytical ultracentrifugation

Analytical ultracentrifugation sedimentation velocity experiments were performed using a Proteome Lab XL-A analytical ultracentrifuge equipped with an An50Ti rotor (Beckman Coulter, Fullerton, CA). The sample (380 μl) and buffer (410 μl) were loaded into each sector of an Epon charcoal-filled two-sector centerpiece. All sedimentation velocity experiments were performed at 25 °C and 42,000 rpm. Absorbance data were collected by scanning the sample cells at intervals of 0.003 cm, monitoring either at 280 nm for protein absorbance or 650nm for Cy5 absorbance. The absorbance signal of protein alone or DNA-protein complexes were maintained between 0.1 and 1.

Continuous sedimentation coefficient distributions, *c*(*s*), were calculated using SEDFIT[48,49]. This analysis yielded individual sedimentation coefficients for each monomer, dimer, and higher oligomer species of proteins as well as a weighted average frictional coefficient (*f/f*_*o*_) for the entire distribution. Calculated sedimentation coefficients were converted to 20 °C water conditions (*s*_*20, w*_) according to:

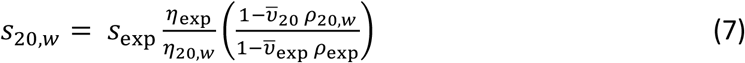

where *ρ*_20, *w*;_ and η_20, *w*_ are the density and viscosity of water at 20 °C, *ρ* _exp_ and η_exp_ are the density and viscosity of the buffer at the experimental temperature of 25 °C, and 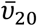 and 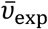 are the partial specific volumes of the protein at 20 °C and at 25 °C. Buffer densities, (*ρ*_exp_) and viscosities (η_exp_) were calculated from buffer composition using SEDNTERP[50]. Partial specific volumes 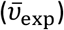 for *Mtb* UvrD1 and Ku were calculated in SEDNTERP using the amino acid composition. Integration of the entire c(s) distribution vs. the integration of an individual sedimentation species was performed and used to calculate the population fraction [48].

For AUC experiments done in the presence of Cy5 labeled DNA, the absorbance signal was collected by scanning the sample cells at 650 nm. Partial specific volumes 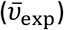 for labeled DNA and the UvrD1-DNA complex were calculated according to:

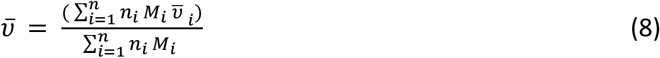

## RESULTS

### Ku does not stimulate UvrD1 dimer-or monomer-catalyzed DNA unwinding on short timescales under single round conditions

We have previously shown that *Mtb* UvrD1 forms a disulfide-bonded dimer under oxidative conditions via a 2B domain cysteine in each subunit and that the dimeric form is an active helicase [42]. The monomeric form of UvrD1 – generated either via reducing conditions or by mutation of cysteine 451 in the 2B domain to alanine – can bind and translocate on single-stranded DNA but lacks helicase activity under previously tested conditions [42]. As it has been reported that Ku stimulates UvrD1 helicase activity [43], we performed DNA unwinding experiments with WT UvrD1 under conditions where dimer formation is favored in the presence and absence of Ku. Specifically, we performed stopped-flow kinetics using a 18bp-dT40 DNA substrate labeled with a Cy5 dye and a black hole quencher (BHQ) on the 5’ and 3’ termini of the blunt end of the duplex respectively. After rapidly mixing a pre-incubated protein-DNA complex with a solution containing 2 mM ATP, 10 mM MgCl_2_, and 5 µM ssDNA trap, we observed an increase in Cy5 fluorescence due to unwinding of the 18 bp duplex. However, contrary to our expectations, we did not observe an increase in either the rate or extent of unwinding in the presence of 800 nM concentrations of Ku **(Figure 1A, red curves)**. We hypothesized that perhaps Ku specifically activates the monomeric form of UvrD1 that we previously observed to completely lack helicase activity under these conditions.

**Figure 1:**
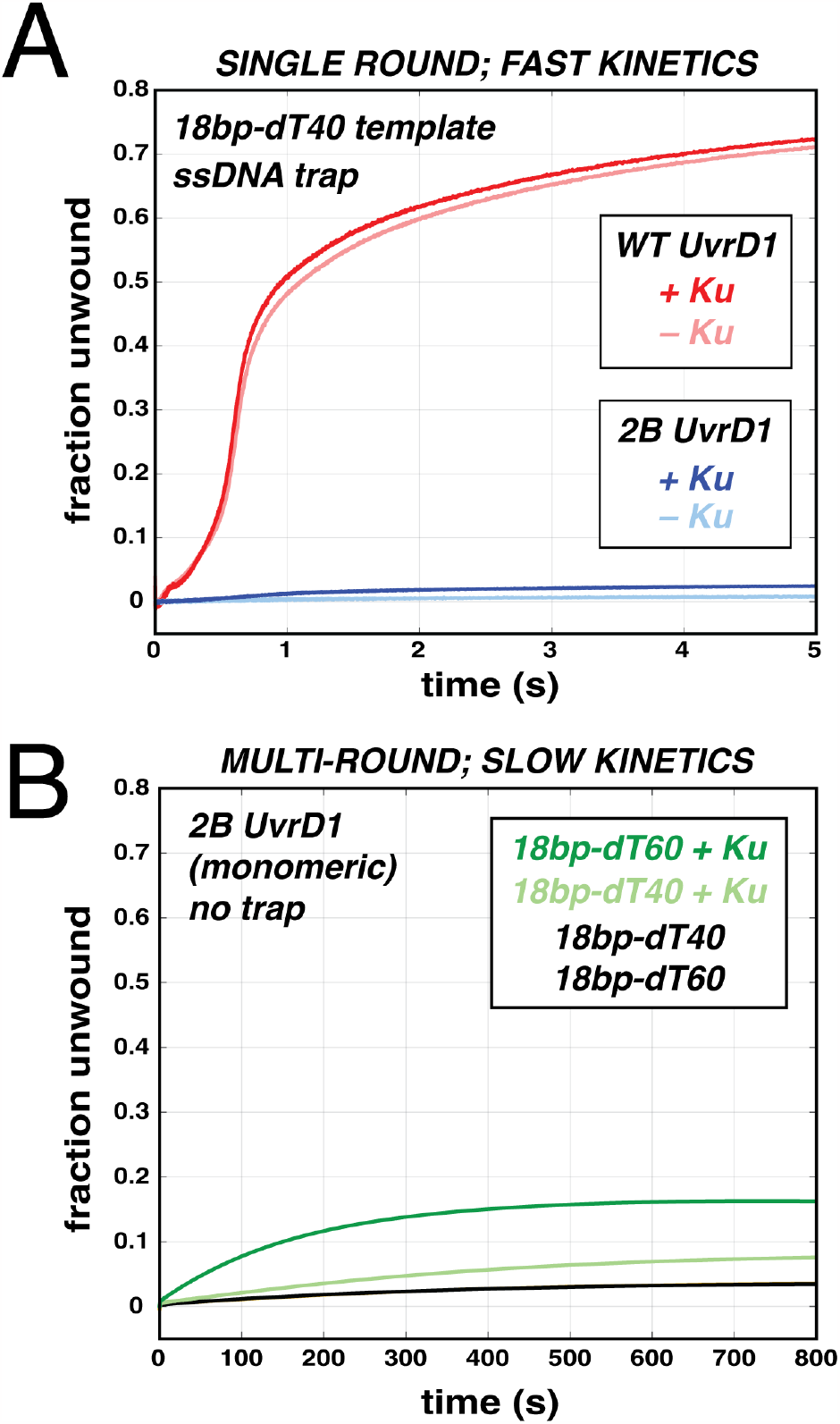
Ku stimulates unwinding of 2BUvrD1 on longer timescales and in multi-round conditions. **(A)** Single-round unwinding of an 18bp-dT40 DNA with WTUvrD1 (red) and 2BUvrD1 (C451A, blue) in the presence and absence of 800 nM Ku at short timescales. (B) Multi-round unwinding of 18 bp duplex with either dT40 or dT60 ssDNA tails with monomeric UvrD1 (2BUvrD1) in the presence (green) or absence (black) of Ku over long timescales. In these experiments, the final concentrations of components are 10 nM DNA substrate, 200 nM UvrD1 and 800 nM Ku.

Stopped-flow experiments were performed with the 2B domain mutant enzyme (C451A, an obligate monomer) to ask whether monomeric UvrD1 could be activated by Ku. Consistent with our previous work, monomer UvrD1 alone displayed no unwinding activity. In addition, we observed no activity upon the addition of 800 nM Ku within 5 seconds **(Figure 1A, blue curves)**. This was true under both single-round and multi-round conditions at these fast timescales **(Supplementary Figure S1A)**. Since the previous Ku-dependent study presented only end point measurements taken after 10 minutes, we next asked whether Ku-dependent activation occurs on longer timescales compared to the rapid unwinding exhibited by dimeric UvrD1 [42].

### Ku activates slow unwinding catalyzed via multiple binding events of UvrD1

Dimeric UvrD1 unwinds 18 bp duplexes with a dT_20_ tail in just over a second **(Figure 1A, red)** [42]. To test the idea that Ku exhibits its effect on longer timescales, we monitored unwinding reactions for over 12 minutes. Consistent with previous observations, DNA unwinding was only observed for UvrD1 dimers under single-round conditions, even at these longer timescales **(Supplementary Figure S1B)**. However, upon the removal of protein trap (ssDNA) to allow multi-round kinetics, a small amount (∼3% after > 13 minutes) of unwinding was observed with the monomeric 2B mutant enzyme **(Figure 1B, black)**. This indicates that this activity is catalyzed by multiple binding events of monomeric UvrD1 in contrast to the processive unwinding displayed by a single dimer. On dT_40_ and dT_60_ tailed duplex substrates, the monomer exhibited very similar activities **(Figure 1B, overlapping black curves)**. We next asked whether Ku might specifically potentiate this activity of the monomer under multi-round conditions. Indeed, in the presence of 800 nM Ku, we observed higher degrees of unwinding and this activation increased with increasing ssDNA tail lengths **(Figure 1B, green curves)**. No DNA unwinding is observed in the presence of Ku alone under these conditions indicating that the effect is due to an enhancement of UvrD1-dependent activity **(Supplementary Figure S2)**.

As would be expected, this Ku dependent activation of monomer unwinding is not unique to the 2B mutant as it is also observed with WT UvrD1 monomers generated by reducing the disulfide bond between the 2B domains with 5 mM DTT **(Figure 2)**. In addition, multi-round unwinding on these time scales can also be observed on longer duplexes (32 bp) and shorter ssDNA tails (20 nt), and the activation increases with increasing Ku concentration **(Figure 2)**. Taken together, these experiments indicate that Ku promotes DNA unwinding by monomers of UvrD1, but only under multi-round conditions over long time scales of 10 minutes.

**Figure 2:**
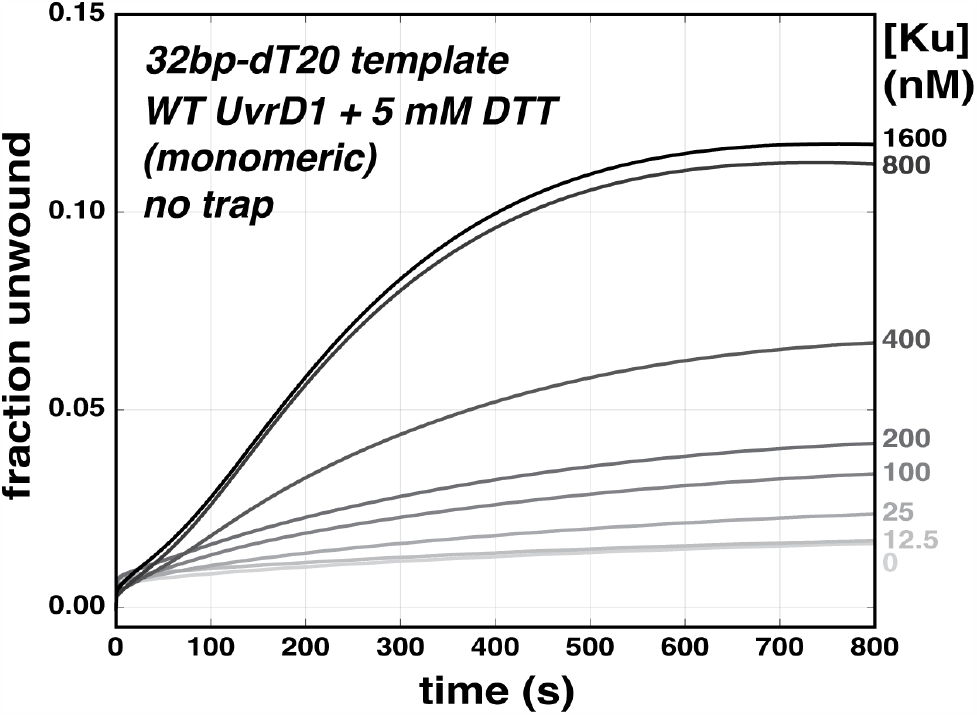
Ku stimulates unwinding by UvrD1 monomers in a concentration dependent manner. Multi-round unwinding of 32bp-dT20 with monomeric WTUvrD1 established by reducing conditions. Increasing Ku concentrations with constant concentrations of UvrD1 monomer (200 nM) and DNA (50 nM) increases unwinding by UvrD1 monomer.

### Ku-dependent activation of full-length unwinding is abrogated in the presence of hybrid traps

The experiments above show that multiple DNA binding events by monomeric UvrD1 are required for DNA unwinding to be observed at long timescales. However, they do not address whether activation requires multiple Ku binding events. We aimed to perform unwinding assays using different traps that would effectively trap Ku. Using a DNA unwinding substrate (32bp-dT_20_) labeled with fluorescein on the 5’ terminus of the blunt end, titration of Ku results in fluorescence quenching **(Figure 3A, black)**. However, Ku does not bind a dT20 ssDNA **(Figure 3A, red)**. These data are consistent with studies of Ku across biology indicating that Ku specifically binds duplex regions of DNA [51,52]. Unwinding assays were thus performed in the presence of UvrD1, Ku, and three distinct nucleic acid traps to assess whether Ku activates UvrD1 in the context of a stably bound Ku-DNA complex or multiple rounds of Ku binding to dsDNA.

**Figure 3:**
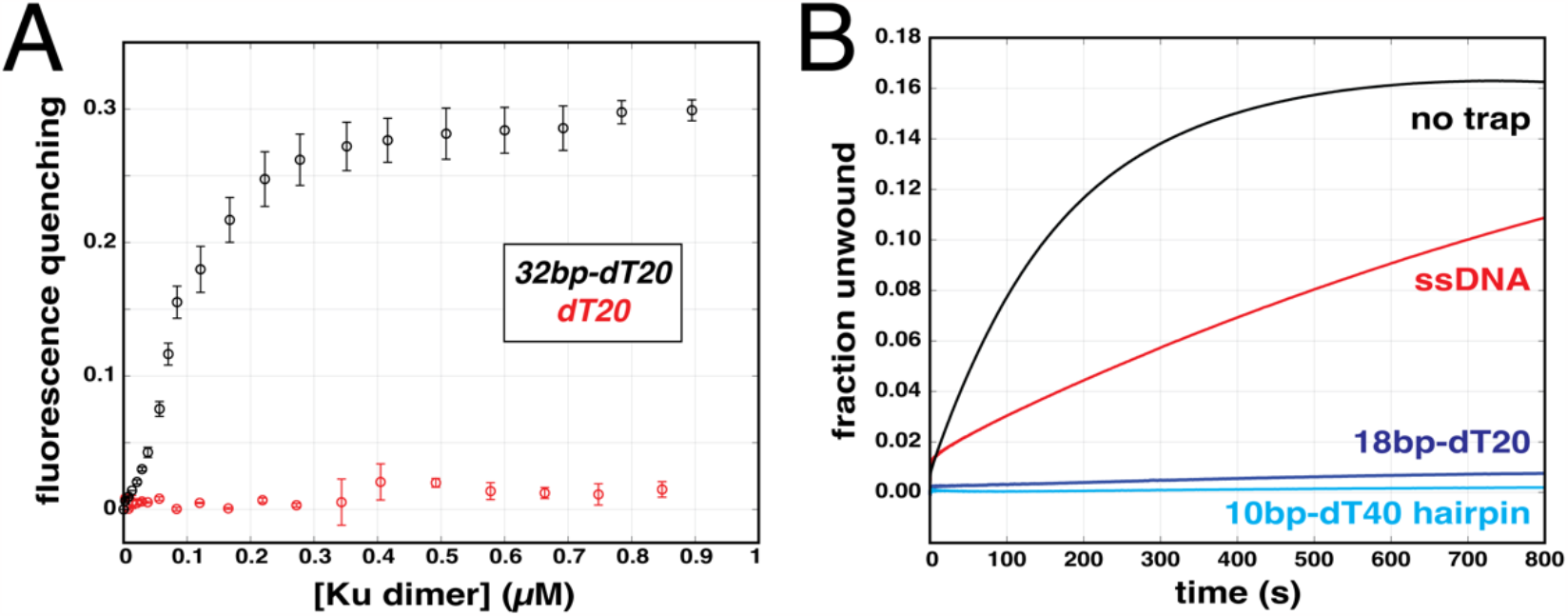
Ku-stimulated UvrD1 monomer unwinding requires multiple Ku binding events. (A) Fluorescence quenching of 50 nM FAM-labeled DNA as a function of Ku dimer concentration (32bp-dT20, black)), and ssDNA (dT20, red). (B) Fraction unwound of a 32bp-dT60 DNA (10 nM) as a function of time in the presence of UvrD1 (200 nM), Ku (400 nM), and no trap (black), ssDNA trap (red), 18bp-dT20 trap (blue), or 10bp-dT40 hairpin trap (cyan) 5µM each.

On 18bp-dT_60_ DNA, hybrid ssDNA/dsDNA traps (18bp-dT_20_ or 10bp-dT_40_ hairpin) resulted in the elimination of DNA unwinding activity **(Figure 3B, blue and cyan)**. In contrast, a ssDNA dT_20_ trap only reduced the kinetics of unwinding **(Figure 3B, red compared to black)**. Given that ssDNA robustly traps UvrD1 alone [42]and Ku does not bind ssDNA alone, these results suggest the existence of a UvrD1-Ku complex, even in the absence of nucleic acid. Furthermore, the ability of traps containing double-stranded regions to abrogate Ku-dependent activation suggests that free Ku dimers must dissociate and rebind the DNA unwinding substrate during the reaction.

### Ku binding to DNA requires magnesium

During our experiments to quantitate the *Mtb* Ku-DNA interaction we discovered a previously unreported requirement of magnesium for stable *Mtb* Ku binding to DNA. The buffer used for unwinding experiments contains 5 mM Mg^+2^ along with 1 mM ATP since both are needed for the helicase activity of UvrD1. However, these constituents were left out in our initial attempts to measure DNA binding of Ku. No Ku-DNA binding was observed using the fluorescence quenching assay in the absence of magnesium **(Figure 4, red)**. However, upon inclusion of 5 mM Mg^+2^ to the binding buffer, robust DNA binding was observed **(Figure 4, black)**. Since Ku is a dimer in solution **(Supplementary Figure S3A)**, we asked whether the presence of Mg^+2^ influenced the distribution of Ku oligomers in the absence of DNA. Analytical sedimentation velocity of 6 μM Ku in the absence and presence of magnesium show that Ku oligomerization does not depend on the presence of Mg^+2^ **(Supplementary Figure S3B)**. Thus, we conclude that magnesium plays a critical role in mediating the interaction between *Mtb* Ku and DNA duplexes. All subsequent binding studies were performed in the presence of magnesium.

**Figure 4:**
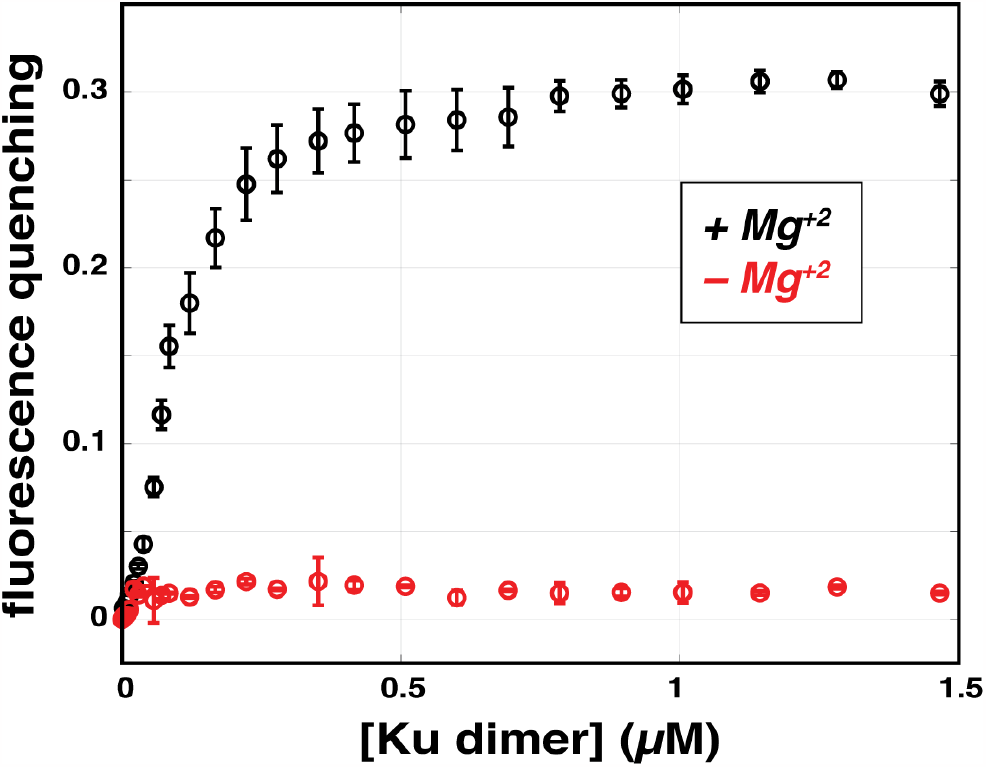
Ku-DNA interaction requires magnesium. Fluorescence quenching of 50 nM 32bp-dT20 DNA labeled with FAM at the blunt end as a function of Ku dimer concentration in the presence (black) and absence (red) of 5 mM magnesium.

### Activation of UvrD1 occurs in the context of multiple Ku dimers bound to the DNA

As part of their role in NHEJ, multiple Ku dimers load at DNA ends and slide inwards along the template [53]. In contrast, *E. coli* UvrD is known to bind ssDNA and has been shown to be positioned at ssDNA/dsDNA junction[54]. To constrain possible models for the interaction between Ku and UvrD1 on the DNA, we wanted to determine whether Ku activation of DNA unwinding was dependent on the number of Ku dimers bound to the DNA substrate. To determine this we examined Ku binding to a 32bp-dT_20_ DNA containg a fluorescein fluorophore attached to the 3’ end of the blunt DNA end. Ku binding to this DNA resulted in a 31% quenching of the fluorescein fluorescence at saturation **(Figure 5A)**. Titration of this DNA with Ku at a DNA concentration of 300 nM showed a linear increase in fluorescence quenching until saturation was reached, indicating that binding is stoichiometric at this high DNA concentration, with a stoichiometry of 2.9 Ku dimers/DNA **(Figure 5A)**, indicating 3 Ku dimers can bind to the DNA at saturation consistent with previous studies suggesting that eukaryotic Ku can bind dsDNA cooperatively and that 3 Ku dimers of *M. smeg* can load on 37bp dsDNA [55,56]. Given the length of the dsDNA region of our DNA unwinding substrate, this suggests an average site size of 10-11 bp, consistent with previous measurements with other Ku systems [57]. Sedimentation velocity in the presence of excess Ku also indicates multiple Ku dimers binding to the DNA unwinding substrate **(Supplementary Figure S4)**.

**Figure 5:**
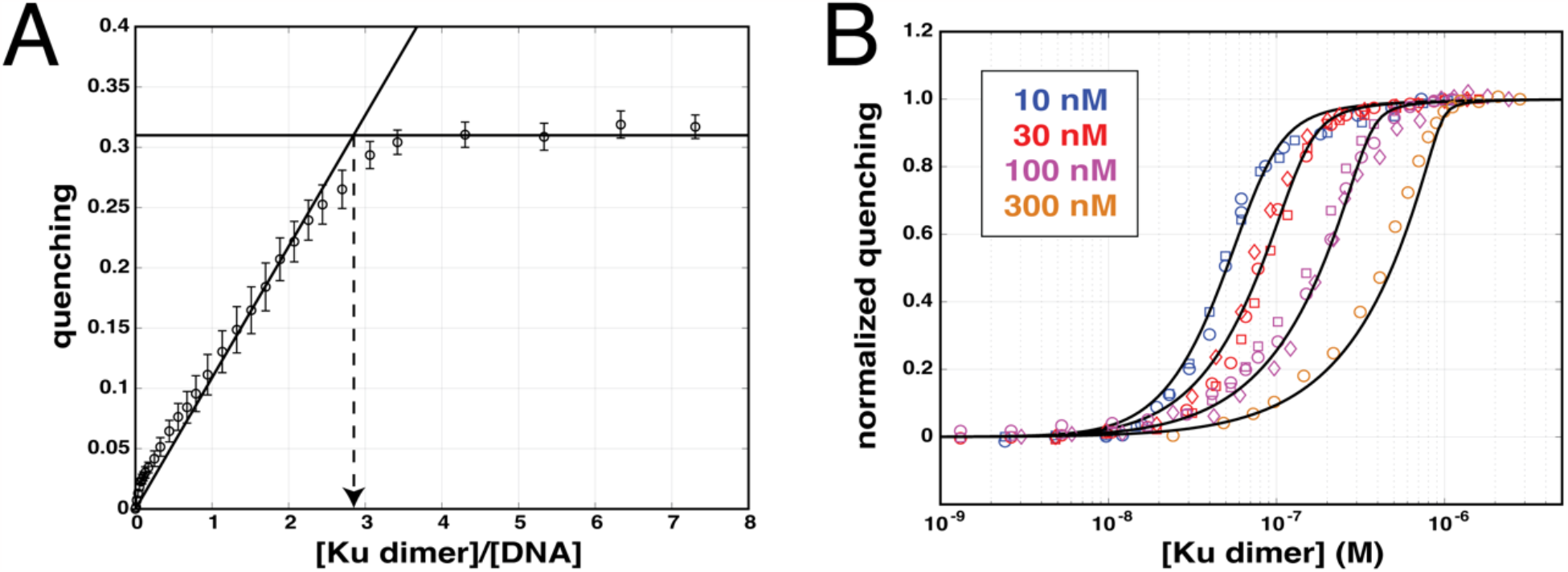
Multiple Ku dimers bind the template with high cooperativity. (A) FAM fluoresence of FAM-labeled 32bp-dT20 DNA is quenched upon titration with Ku. Results of a titration of 300 nM DNA with Ku under stoichiometric binding conditions in TRIS pH 8.0, 75mM NaCl, 20% glycerol amd 5mM MgCl2 yields a stoichiometry of 3 Ku dimers per DNA at saturation (dotted line). (B) Normalized fluorescence quenching as a function of Ku dimer concentration with 10, 30, 100, and 300 nM DNA substrate. Fits from the model described in the text are shown as solid lines.

To obtain quantitative information about Ku binding affinity and cooperativity, we performed titrations at multiple DNA concentrations of 10, 30, 100, and 300 nM. The results are shown in **Figure 5B**, plotted as fluorescence quenching as a function of total Ku dimer concentration. To analyze these titrations quantitatively, we first performed a binding density function analysis [46] to determine the relationship between the average extent of Ku binding per DNA and the average fluorescence quenching. This approach relies on the fact that free ligand (Ku dimer) concentration determines the average extent of Ku binding independent of DNA concentration. The average fluorescence quenching signal showed a linear dependence on the average extent of Ku bound until saturation at a stoichiometry of ∼2.5 Ku dimers bound per DNA **(Supplementary Figure 5)**, consistent with the estimate from the stoichiometric titration **(Figure 5a)**.

We next used a nearest-neighbor cooperative model for non-specific binding of a large ligand (Ku dimer) to a finite linear lattice (DNA substrate) [47] (see Materials and Methods for details). This model assumes Ku dimers bind to the duplex DNA with equilibrium association constant K and with nearest neighbour cooperativity indicated by the cooperativity parameter, ω. We assumed an occluded site size, n=11 bp per Ku dimer, as estimated from the stoichometric titration data, and a duplex DNA lattice length of 33 bp (one more than the actual duplex DNA length of 32 bp). The binding polynomial, P, for this model, using the free DNA as reference state, is given by Eq. (1), and the average extent of Ku dimer bound per DNA, <Ku>, and is given in Eq. (2).

This model has only three parameters, K, ω, and Q_max_, which can be estimated from a global non-linear least squares (NLLS) fit of the 4 titrations shown in **Figure 5B**. The best fit parameters obtained from the NLLS fit are **K = (3.3 ± 1.8) × 10**^**4**^ **M**^**-1**^ **and ω = (2.3 ± 1.9) × 10**^**4**^. The solid curves in **Figure 5B** are simulations using Eq. (2) and these best fit values of K and ω. Since a non-cooperative binding system would have a value of ω = 1, this indicates that Ku binds with quite high positive cooperativity to duplex DNA. The large uncertainties associated with the estimates of K and ω result from the highly cooperative nature of the Ku-DNA binding, such that the values of K and ω are highly anti-correlated. **Supplementary Figure 6** shows that constraining ω = 1, does not provide a good fit to these titrations, again indicating that Ku dimers bind the DNA substrate with high cooperativity.

**Figure 6:**
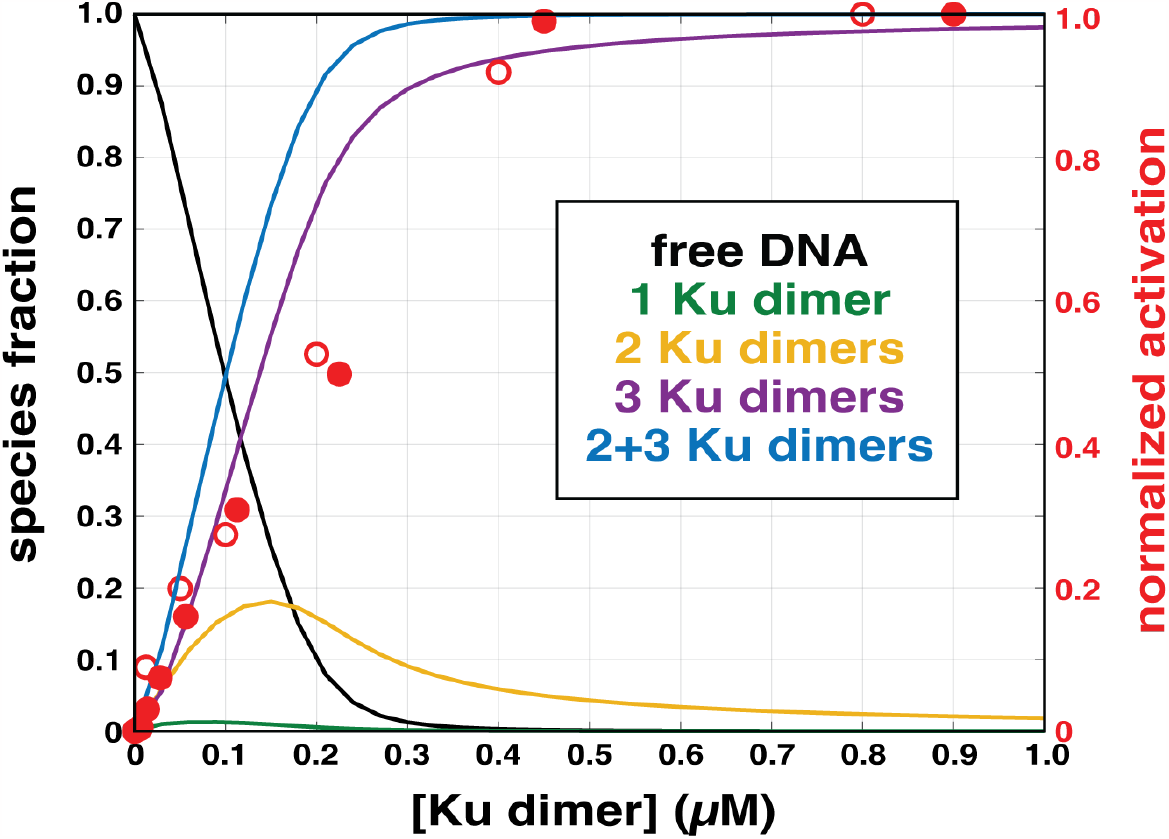
Helicase activation occurs in the presence of 3 Ku dimers bound. DNA-bound species fractions as a function of total Ku dimer concentration in the presence of 50 nM DNA substrate shown along with the fold-change of DNA unwinding activation using the same DNA concentration. The open and closed circles are activation data from two separate experiments.

Figure 6. shows a plot of the species fraction of DNA bound with one, two or three Ku dimers as a function of total Ku dimer concentration at 50 nM DNA calculated using Eqs. (3-5). These distributions, calculated using K = 3.3 × 10^4^ and ω = 2.3 × 10^4^, are compared with the Ku-concentration dependent activation curve obtained under the same DNA concentration. This comparison shows that UvrD1 monomer activation can occur in the context of three Ku dimers bound to the DNA substrate. If the three Ku bound DNA were inhibitory, then the activation curve would increase to a peak and then decrease, as indicated for the one and two Ku bound DNA species in **(Figure 6, yellow and green)**. Thus, the triply-bound state is not inhibitory and, instead, is responsible for the majority of the activation of UvrD1 unwinding. Due to the high cooperativity of the system, the data do not allow us to evaluate the relative abilities of the other distinct Ku-bound structures to activate. For example, summing the doubly-bound and triply-bound species fractions closely follows that of the triply bound alone **(Figure 6, blue)**. As such, it remains to be seen whether a single Ku dimer or two Ku dimers can activate as efficiently as the fully saturated DNA substrate.

### *Mtb* Ku can bind DNA with single-strand 3’-dT_20_ overhangs

Our *in vitro* DNA unwinding substrate has two ends: a single-stranded end with which UvrD1 interacts and a double-stranded blunt end with which Ku may interact to load on the DNA. Previous studies have suggested that *Mtb* Ku can also load onto DNA with ssDNA overhangs [57]. To test this idea and investigate the determinants of *Mtb* Ku DNA binding further, we performed sedimentation velocity experiments on Ku in the presence of fluorescently labeled DNA with different end types. As expected, no Ku-ssDNA complexes were observed with 1 μM ssDNA (dT_20_) and 8 μM Ku **(Figure 7A)** and Ku-dsDNA complexes were observed with blunt-ended 50 bp duplex fluorescently labeled internally to control for any potential fluorophore effects on Ku binding/loading at the DNA end **(Figure 7B, green)**. The complexes observed on the dsDNA show a broad distribution of sedimentation coeffficients consistent with multiple Ku dimers bound to the DNA **(Figure 7B, green, inset)**. In the presence of a similarly internally labeled 50 bp dsDNA with a 3’-dT_20_ single-stranded tail on each end, a similar distribution of protein-DNA complexes was observed **(Figure 7B, red, inset)**. These data show that *Mtb* Ku can bypass single-stranded DNA tails of at least 20 bases and bind internal dsDNA duplex regions and suggests that, in the cell, *Mtb* Ku may load on free DNA ends regardless of the overhang status.

**Figure 7:**
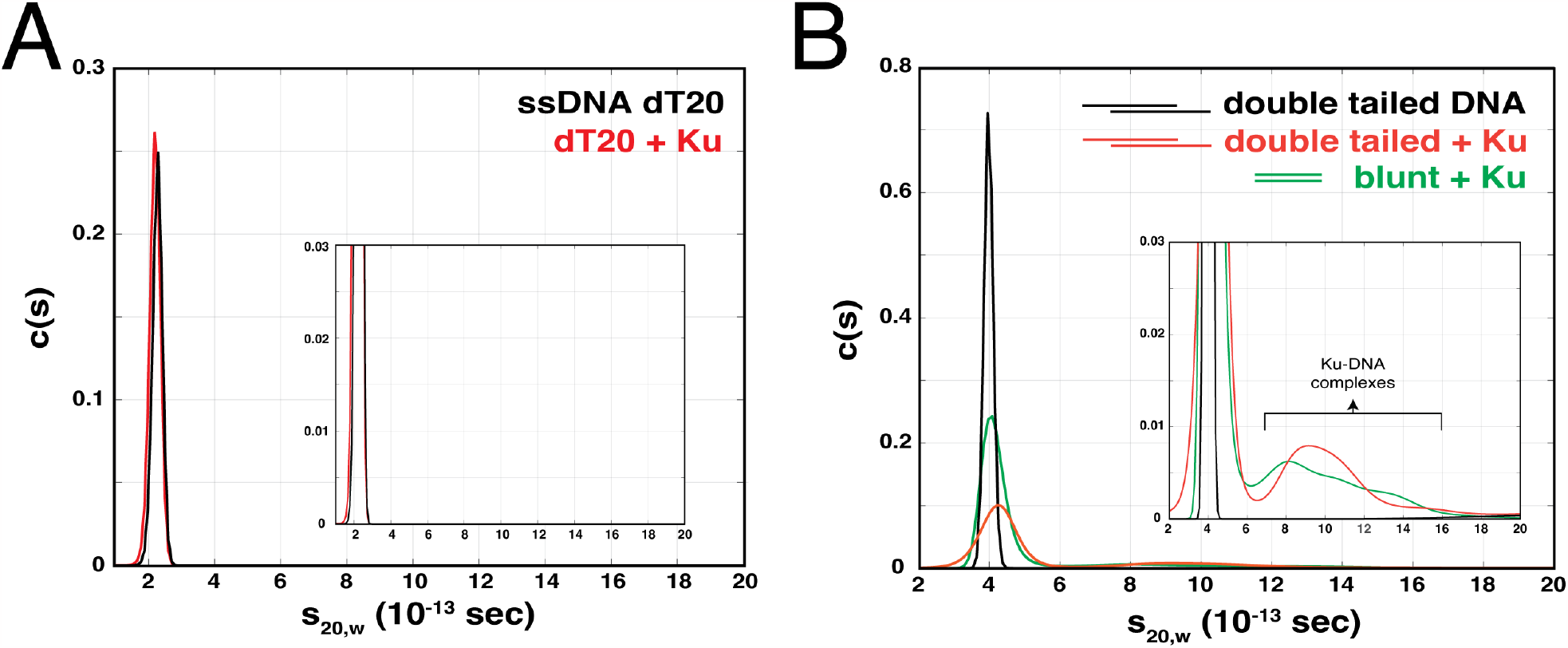
Mtb Ku binds DNA with double dT20 single-strand 3’-overhangs. **(A)** AUC sedimentation velocity of dT20 ssDNA labeled with Cy5 at 5’ end alone (black) 1 μM and in the presence of 8 μM Ku (red). **(B)** AUC sedimentation velocity of 50 bp dsDNA internally labeled with Cy5 with two dT20 single-stranded tails on either end alone (black) 1 μM and in the presence of 8 µM Ku (red). The addition of Ku to blunt (50 bp internally labeled with Cy5 as control shown in green).

### The C-terminal Tudor domain of UvrD1 is required for the activation of unwinding

Previous studies described an interaction between Ku and the C-terminal region of *Mtb* UvrD1 via yeast-two hybrid assays[26]. Further work using *M. smegmatis* UvrD1 showed that deleting the C-terminal 91 amino acids from UvrD1 prevented its interaction with Ku [44]. However, the same truncated construct maintained the ability to be activated by Ku in gel-based DNA unwinding assays. One caveat to the interpretation of these experiments is that they lacked the ability to separate dimer and monomer UvrD1 effects. Given our identification of the UvrD1 monomer as the activated oligomer by Ku under multi-round conditions, we returned to these observations and asked whether the C-terminal region of UvrD1 was required for Ku-dependent helicase activation.

Based on alignments with other UvrD-family members including *M. smegmatis* UvrD1, we deleted the C-terminal 46 amino acids of *Mtb* UvrD1 which make up the conserved C-terminal Tudor domain **(Supplementary Figure S7)** [58]. We first characterized the effect of this UvrD1ΔTudor mutant on the interaction between UvrD1 and the DNA substrate itself. Sedimentation velocity with Cy5-labeled DNA unwinding substrate revealed that both monomers and dimers of UvrD1ΔTudor bind DNA in about the same ratio as WT **(Supplementary Figure S8)**. In fact, at the same concentrations, more monomers and dimers of UvrD1ΔTudor were bound to DNA, suggesting a higher affinity. Furthermore, UvrD1ΔTudor in the presence of DTT is monomeric like WT UvrD1 **(Supplementary Figure S9**) and shows increased DNA unwinding as compared to WT UvrD1 suggesting that the Tudor domain may be inhibitory under these conditions **(Figure 8, cyan)**. However, DNA unwinding assays performed with reduced UvrD1ΔTudor in the presence and absence of Ku revealed a lack of activation **(Figure 8, purple and cyan)** showing that Ku activation is Tudor domain dependent.

**Figure 8:**
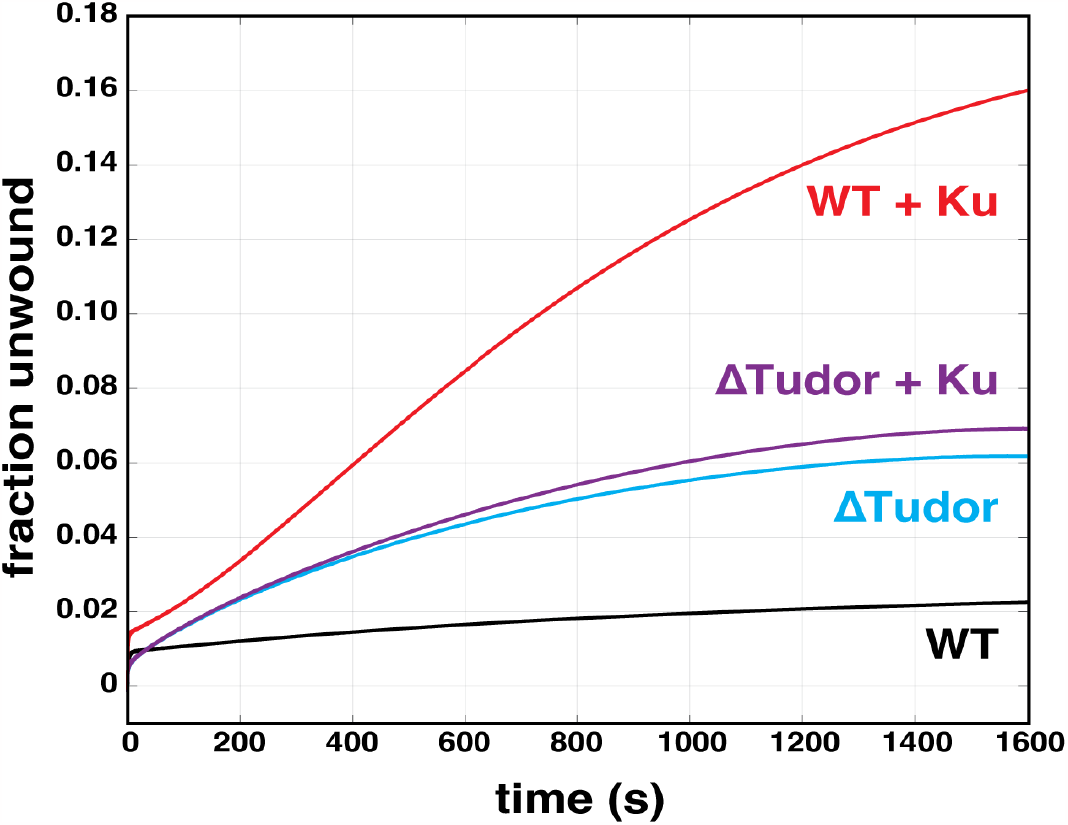
The C-terminal Tudor domain of UvrD1 is required for Ku-dependent activation of monomer unwinding. **(A)** The addition of Ku activates UvrD1 monomer unwinding in reducing conditions (red and black). UvrD1Δ1Tudor shows enhanced unwinding relative to WT (cyan and black) 200nM, but the addition of 400 nM Ku dimer does not have any effect (purple). Each trace shown was performed on 10 nM 32bp-dT20 unwinding substrate.

## DISCUSSION

Processive DNA helicases use energy from ATP hydrolysis to translocate on single-stranded DNA and unwind double-stranded DNA [[59,60]. Extensive biochemical studies have shown that UvrD-like helicases unwind DNA processively as homodimers in the absence of accessory factors[31,39,61]. We have previously shown this to be true for *Mtb* UvrD1[42] where dimerization of UvrD1 results in processive helicase activity and is linked to redox potential through the oxidative formation of a disulfide bond between 2B domains **(Figure 9, top panel)**. Mutagenic or chemical disruption of the disulfide bond resulted in monomeric UvrD1 which did not possess helicase activity under single-turnover conditions [42].

**Figure 9:**
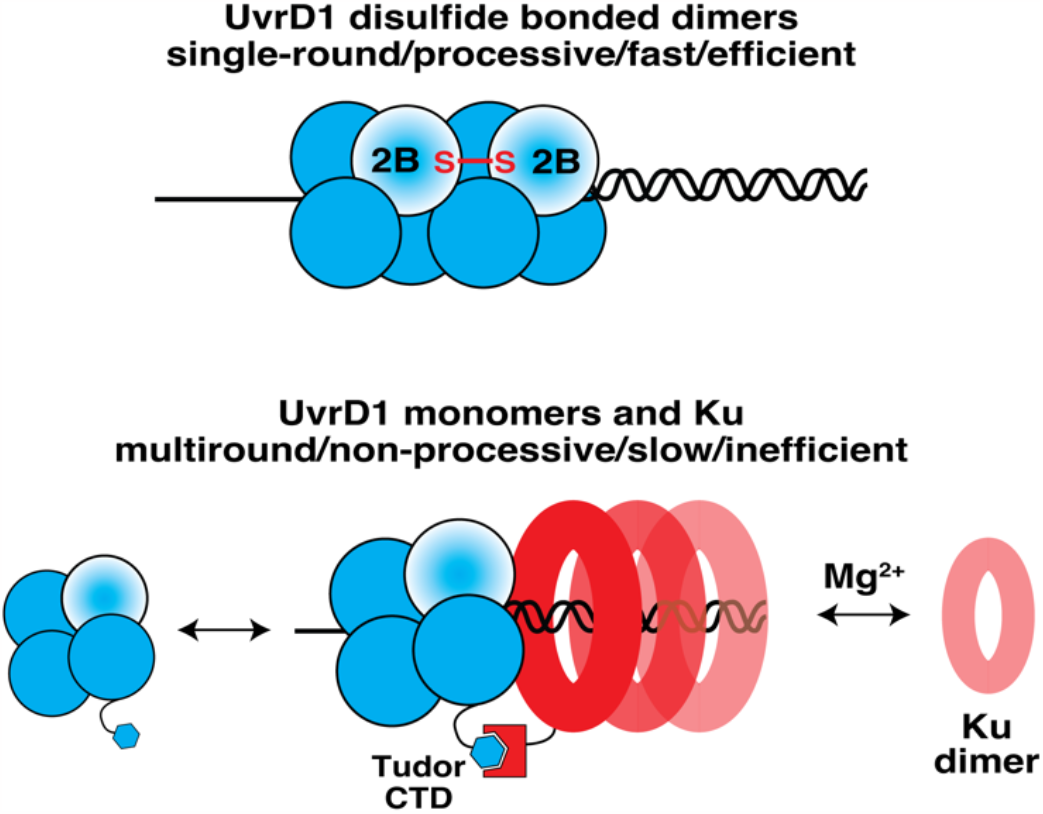
Models for UvrD1 unwinding mechanisms. **(top)** A disulfide bond between the 2B domain of two UvrD1 molecules leads to the activation of processive and rapid helicase activity[42]. **(bottom)** Monomers of UvrD1 exhibit a measurable amount of multi-round unwinding activity at long time scales which is activated by the presence of multiple Ku dimers on the double stranded DNA substrate.

However, there are two fundamentally different ways to assay for helicase activity. Single-round experiments probe whether an enzyme (or enzyme complex) can productively unwind a DNA substrate of a particular length in the context of a single binding event. To do so, the enzyme must possess the ability to take processive steps along the DNA equal to the length of the duplex region without dissociating. If enzyme does dissociate from the DNA, a suitable enzyme trap prevents re-binding. In contrast, multi-round experiments, performed in the absence of a trap, allow for dissociated enzymes to re-bind. Under these conditions, the enzyme does not need to possess a processivity on the order of the length of the DNA. Here, we show that although the monomeric form of UvrD1 does not display single-round DNA unwinding activity even on the shortest DNA substrates, it does exhibit a small (*i*.*e*., 3%) degree of DNA unwinding under multi-round conditions which is activated by *Mtb* Ku. This activity is approximately 100 times slower than that exhibited by the dimer. In fact, the two timescales for DNA unwinding are clearly observed in a single experiment when a mixture of dimers and monomers are present **(Supplementary Figure S10)**.

Previous work by others showed that *Mtb* Ku, an accessory factor involved in NHEJ, stimulates the helicase activity of UvrD1 [43,44]. In other examples of UvrD-family enzyme activation, the effects of auxiliary factors could be seen on monomers, dimers, or both species [62–64]. Considering this, we asked which oligomeric species of UvrD1 was involved in the Ku-dependent mechanism. Since UvrD1 can be readily converted to monomers by the addition of a reducing agent or by single point mutation of cysteine to alanine at position 451 in the 2B domain, we could directly test whether Ku enhances the helicase activity of monomers, dimers, or both. We found that Ku did not affect unwinding by the processive dimer of UvrD1 but increased unwinding by UvrD1 monomers about five-fold. This stimulation of unwinding is only observed over long timescales of 10 minutes in multi-round conditions and in the absence of hybrid ssDNA-dsDNA traps. This is consistent with previous studies showing Ku-stimulated unwinding on longer time scales using excess single-stranded DNA traps [43].

Although published studies of Ku-stimulated unwinding by UvrD1 report 80-90% DNA unwound [43], we observed a maximum of only 15% DNA unwound under monomeric UvrD1 conditions. This discrepancy is likely due to the absence of reducing agent in the previous work that would lead to unwinding by both dimers and monomers. When a mixture of dimers and monomers exist, we also observe 80% of the DNA unwound **(Supplementary Figure S10)**. In addition, previous studies were performed at higher temperatures. When we performed a comparison of the unwinding kinetics at 37 °C compared to 25 °C, we observed accelerated rates and a 50% increase in the fraction unwound of a dT_20_-18 bp substrate **(Supplementary Figure S11)**. [65]

All previous studies investigating the binding of Ku proteins to DNA were performed in presence of Mg^+2^ [55,66]. In the process of our studies presented here, we unexpectedly found that the interaction of *Mtb* Ku and DNA requires the presence of magnesium. We have not explored the concentration dependence or stoichiometry of this requirement, nor have we investigated the ability of other divalent cations to substitute for magnesium. These aspects of the Ku-DNA interaction would be interesting for future studies.

Our AUC sedimentation velocity data show that *Mtb* Ku exists as a dimer in solution and can bind to blunt ended dsDNA, consistent with previous studies [67,68]. We also showed that a single-strand overhang does not prevent the loading of *Mtb* Ku on dsDNA and that multiple Ku dimers coat our DNA substrates. Taken together, our data suggest a model where activation occurs in the context of a UvrD1 monomer bound to the single-stranded side of the junction interacting with a Ku dimer bound to the double-stranded side of the junction. Since activation increases as more Ku dimers are loaded onto the DNA, we hypothesize that the loading of additional Ku’s serves to increase the probability of a junction proximal Ku by restricting the available space on the duplex region of the DNA **(Figure 9, bottom panel)**.

On a 18bp-dT_20_ substrate, this multi-round activity of Ku-activated monomers leads to unwinding of only about 5% of the available DNA over the course of tens of minutes. The lengthening of a flanking 3’-single-stranded DNA tail on the duplex DNA leads to more efficient unwinding over similar timescales **(Supplementary Figure S12)**. This observation suggests that either the rate of binding of UvrD1 monomer to the DNA limits the yield of the reaction and/or that the ability to bind multiple monomers simultaneously on a DNA substrate increases the efficiency of the reaction. Future experiments are required to investigate this effect further.

It is not clear which specific DNA repair pathways might utilize the interaction between Ku and UvrD1. Double-stranded breaks are processed either by resection-independent or resection-dependent mechanisms in mycobacteria [10]. The canonical NHEJ pathway utilizes Ku-LigD complexes to seal breaks without single-stranded resection. In addition to ligase activity, the mycobacterial version of LigD has both nuclease and polymerase activities which aid in the trimming of overhanging flaps and the filling in of nucleotides prior to ligation [15]. The resection-dependent pathways require the production of single-stranded DNA overhanging regions to be used either for homology searches (i.e., homologous recombination or gene conversion) or for single-stranded annealing (SSA) [9,69]. These two pathways depend on the combined helicase/nuclease activities of AdnAB and RecBCD respectively and are independent of Ku [70]. In these contexts, one possible hypothesis based on our data is that UvrD1 controls the timing of and partitioning between these different strategies. Its ability to interact with Ku may inhibit NHEJ, while multiple rounds of binding and unwinding may prepare the substrate for resection by other enzymes. This would have the effect of funneling the repair away from non-homologous end joining and into homology-based mechanisms. The concentration and oligomeric status of UvrD1 would then provide a means for the regulation of fluxes in each pathway.

SF1 family helicase monomers have previously been shown to be activated through protein-protein interactions with accessory factors. For instance, *E. coli* UvrD monomer and dimer helicase activities are both stimulated by MutL, and *E. coli* Rep monomer and dimer can be activated by PriC [62–64]. In contrast, we distinctly did not observe a Ku-dependent increase in unwinding by the UvrD1 dimer **(Supplementary Figure S1A)**. We note that perhaps the presence of the covalent dimer, absent in the case of *E. coli* UvrD and Rep, contributes to this difference.

The activation of *E. coli* Rep monomer helicase activity by PriC depends on the C-terminal domain and previous yeast-two-hybrid results suggest that the CTD of *Mtb* UvrD1 interacts with the N-terminus of Ku [43]. Previous studies have shown that removing the entire C-terminal domain of *M. smegmatis* UvrD1 including the disordered linker and the Tudor domain leads to abrogation of super shifted Ku-UvrD1-DNA complex in a native gel assay, but Ku can still stimulate DNA unwinding by a similar construct of *Mtb* UvrD1[26]. We removed the C-terminal Tudor domain (46 amino acids) of *Mtb* UvrD1 to test directly if the Tudor domain of *Mtb* UvrD1 played a role in the Ku-dependent activation and found that this increases the amplitude of multi-round unwinding by UvrD1 both in the absence and presence of DTT **(Supplementary Figure S13)**, suggesting that the Tudor domain itself is autoinhibitory for DNA unwinding. The observation is reminiscent of the finding that removal of the autoinhibitory 2B domain leads to activation of monomeric Rep helicase activity [71]. Furthermore, although basal levels of DNA unwinding were increased, the ΔTudor construct was no longer activated by Ku **(Figure 8)**. This observation is consistent with the physical interaction of the Tudor domain with Ku and supports a hypothesis that this interaction eliminates the autoinhibitory effect of the Tudor domain on DNA unwinding.

The results reported here should aid our understanding of the physiological role of UvrD1. Specifically, one could dissect the roles of the UvrD1 dimer and Ku-UvrD1 monomer complex *in vivo* by replacing WT UvrD1 with either the 2B cysteine mutant which is an obligate monomer or with the UvrD1ΔTudor construct which can still form dimers but can no longer interact with Ku. Additionally, one might test the hypothesis that the UvrD1 dimer is active in NER while the Ku-UvrD1 monomer complex participates in alternative pathways of repair including DSBR.

The potential inhibitory role of the Tudor domain is particularly interesting given that the same domain has been shown to underlie the interaction of *E. coli* UvrD and *B. subtilis* PcrA with the RNA polymerase beta-subunit during transcription-coupled NER (TCR) [40,72,73]. In TCR, lesions on the template strand cause RNAP pausing during elongation. This paused state becomes a substrate for either UvrD or Mfd, which then process the complex to both remove the polymerase from the lesion by either backtracking or forward tracking respectively and recruit downstream NER factors [74–77]. In the case of UvrD, it has been suggested that a dimer of UvrD is needed to produce backtracking [74,78]. The Tudor domain results here are likely extendible to homologous systems as we observed that *Mtb* Ku also activates *E. coli* UvrD that contains a Tudor domain, but not *E. coli* Rep that does not contain a Tudor domain **(Supplementary Figure S14)**. This possibility would also suggest that the Tudor domain may serve as a protein-protein interaction hub for UvrD1 and that the partitioning of UvrD1 into various repair pathways would ultimately be controlled via competition for this domain.

## AUTHOR CONTRIBUTIONS

A.C. and E.A.G. designed research; A.C. performed experiments and analyzed the data. B.N. provided *E. coli* UvrD and Rep and assisted with data analysis. A.K. analyzed the Ku-DNA titrations. A.C., T.M.L. and E.A.G. wrote the paper. T.M.L. and E.A.G. guided the project.

## ACKNOWLEDGEMENTS

We thank Rahul Chadda from Edwin Antony’s lab for their help in using their fluorimeter instrument for preliminary anisotropy assays. We thank Galburt lab members for their comments on the manuscript.

