## Supplemental Information for "*Mycobacterium tuberculosis* Ku stimulates multi-round DNA unwinding by UvrD1 monomers"

Ankita Chadda[1], Alexander G. Kozlov[1], Binh Nguyen[1],
Timothy M. Lohman[1]* and Eric A. Galburt[1]*
[1]Department of Biochemistry and Molecular Biophysics, Washington University School of Medicine, St. Louis, MO, 63110, USA.

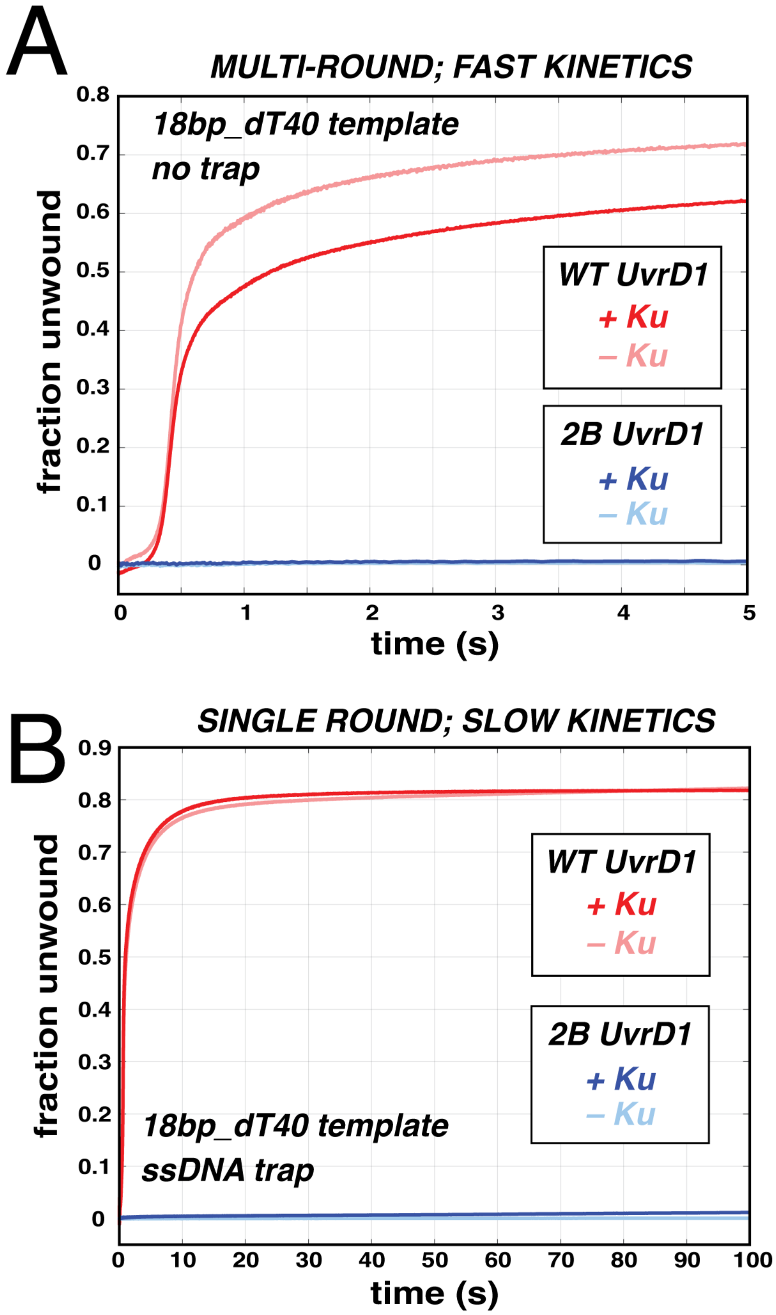


***Figure S1: DNA unwinding by UvrD1 dimers is not stimulated by Ku on short time scales in multi-round conditions nor in single-round conditions at longer timescales.*** **(A)** Multi-round DNA unwinding by WT UvrD1 in the absence of DTT where it's a mixture of monomers and dimers in the presence and absence of Ku (red traces). Multi-round DNA unwinding by 2B UvrD1 where removal of cysteine 451 leads to abrogation of dimerization in the presence and absence of Ku (blue traces). **(B)** Ku did not affect single-round unwinding by WT UvrD1 in the absence of DTT at longer timescales (red traces). Nor did Ku affect single-round unwinding by 2B UvrD1 under these conditions (blue traces).

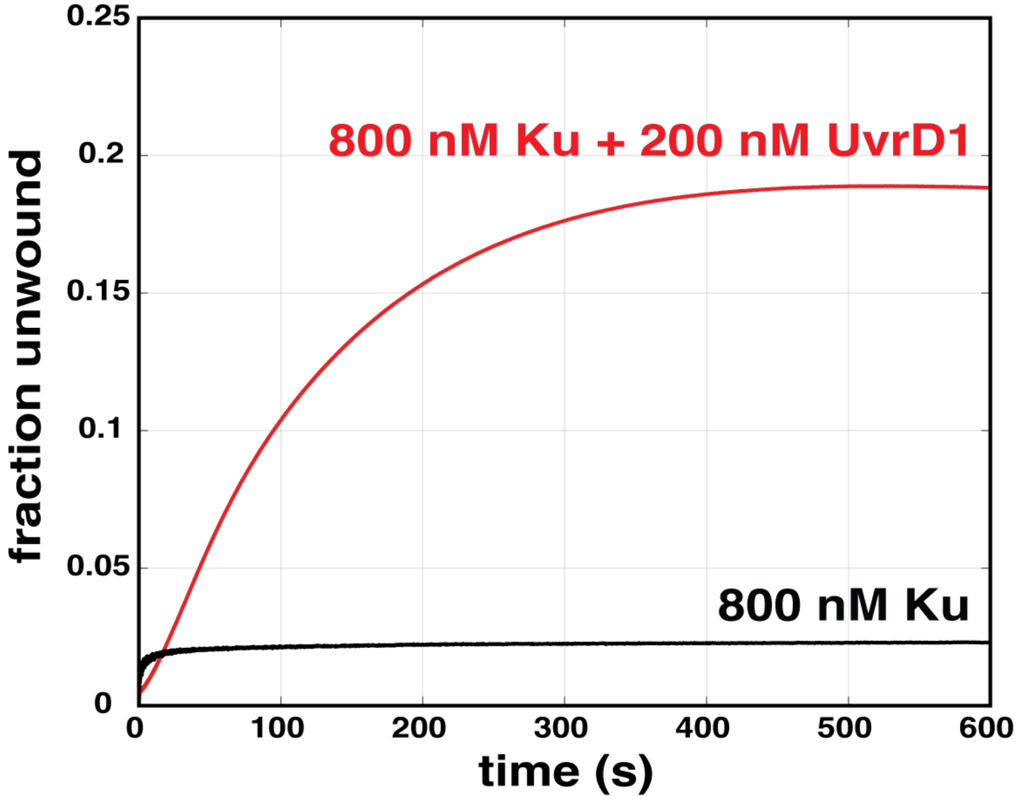


***Figure S2: Ku alone does not account for the DNA unwinding observed in the presence of UvrD1***. DNA unwinding traces are shown with 800 nM Ku in the presence (red) and absence (black) of 200 nM UvrD1 monomers.

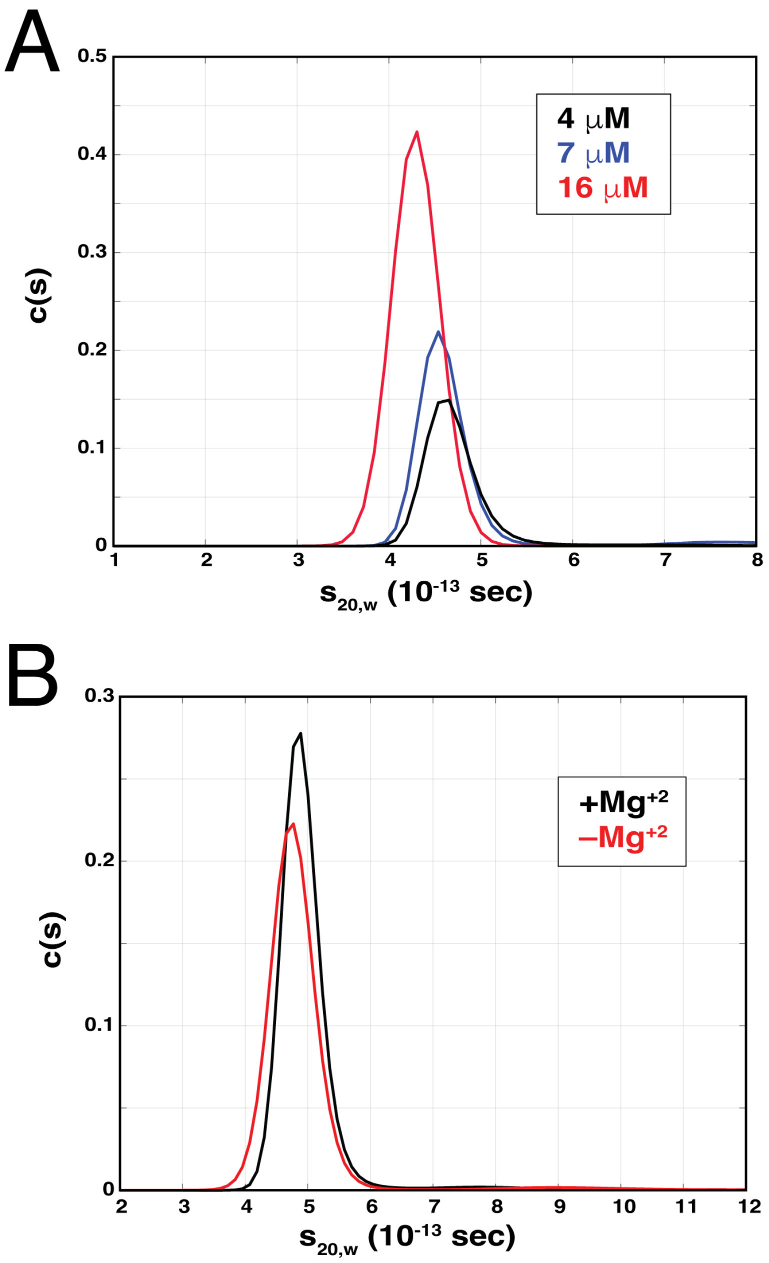


***Figure S3: Mtb Ku is a dimer in the presence and absence of magnesium. (A)*** Ku was run in AUC sedimentation velocity at the three concentrations indicated and, in each case, the data suggested a molecular species consistent with that of a dimer. ***(B)*** 3 μM Ku was run in AUC sedimentation velocity in the presence (black) and absence (red) of 5 mM $MgCl_2$. The data are consistent with a model where magnesium has no effect on the oligomeric state of Ku.

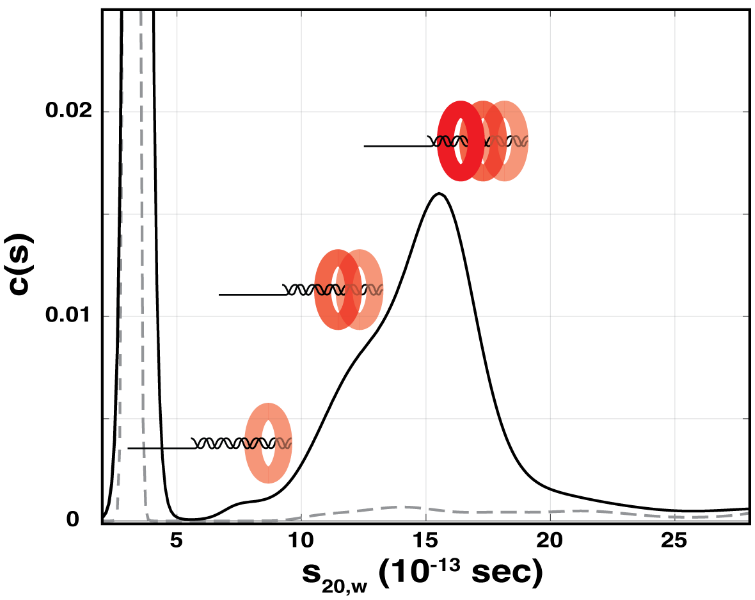


***Figure S4: Sedimentation velocity profile showing multiple Ku-DNA structures.*** 1 μM labeled 32bp-dT20 substrate alone (dashed line) in the presence of 4 μM Ku dimer (solid line). Singly, doubly, and triply bound peaks are indicated.

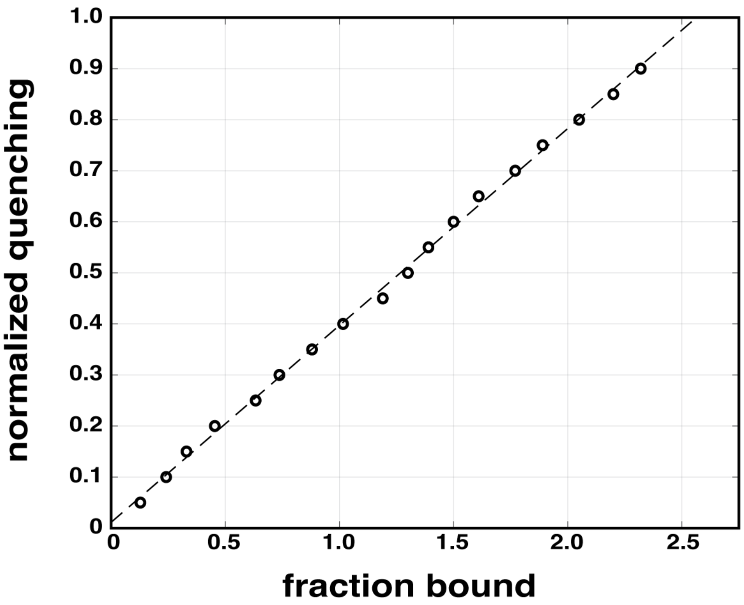


***Figure S5: Normalized quenching as a function of average fraction bound.*** The quenching signal is linearly dependent on the average fraction bound and suggests 2.5 Ku dimers bound per DNA.

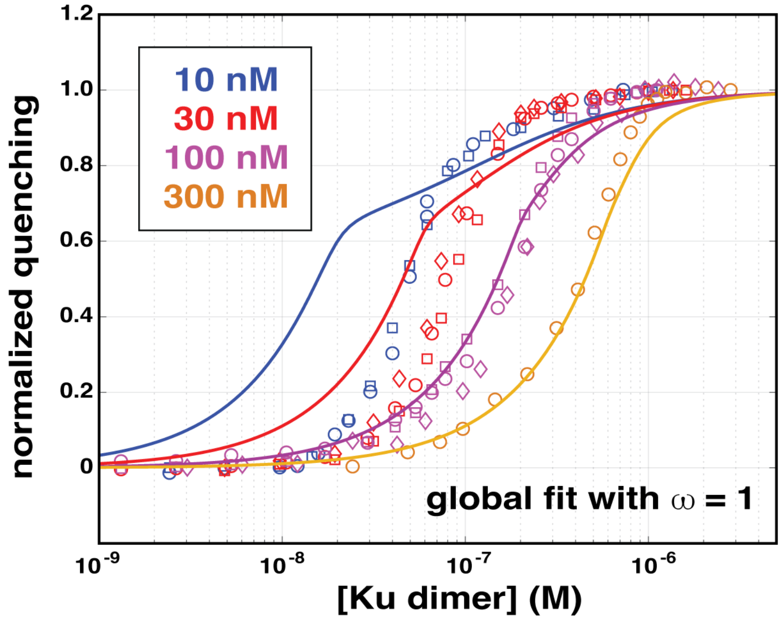


***Figure S6: A global fit of the Ku titration data set with a model in the absence of cooperativity.*** The normalized quenching data from Ku titrations at different concentrations of DNA substrate (10, 30, 100, and 300 nM) are shown. A non-cooperative model was unable to fit the data (solid lines, compare with main text Figure 5).

```
E.coli_UvrD                        QKLTLTYAETRRLYGKEVYHRPSRFIGELPEECVEEVRLRATVSRPVS------------     655
B.stearothermophilus_PcrA          EELVLTSAQMRTLFGNIQMDPPSRFLNEIPAHLLETASRRQA------------------     654
Mtb_UvrD1                          QRLYVSRAIVRSSWGQPMLNPESRFLREIPQELIDWRRT-APKPSFSAPV------SGAG     699
M.smeg_UvrD1                       QRLYLSRAKVRSSWGQPMLNPESRFLREIPQELIDWRRIEAPTPAYAAPGRMSSSGGGGI     710
                                   :.* :: *  *  :*:   .  ***: *:* . ::

E.coli_UvrD                        --------HQRMGTPMVENDSGYKLGQRVRHAKFGEGTIVNMEGSGEHSRLQVAFQGQ-G     706
B.stearothermophilus_PcrA          --GASRPAVSRPQA--SGAVGSWKVGDRANHRKWGIGTVVSVRGGGDDQELDIAFPSPIG     710
Mtb_UvrD1                          RFGSARPSPTRSGA-SRRPLLVLQVGDRVTHDKYGLGRVEEVSGVGESAMSLIDFGSS-G     757
M.smeg_UvrD1                       GFGSPRPSPNRPGGGRNKPLMVLQPGDRVTHDKYGLGRVEEVAGTGESAMSLIDFGSA-G     769
                                             *            : *:*. * *:* * : .: * *:     : * .  *

E.coli_UvrD                        IKWLVAAYARLESV     720
B.stearothermophilus_PcrA          IKRLLAKFAPIEKV     724
Mtb_UvrD1                          RVKLMHNHAPVTKL     771
M.smeg_UvrD1                       RVKLMHNHAPLQKL     783
                                      *:  .* :.:
```

deleted C-terminal
sequences in this work and others
(see references X, X, X)

Tudor domain sequence based
on Sanders, et al. 2017
doi: 10.1093/nar/gkx074

***Figure S7: Sequence alignment of C-terminal domain of SF1 helicases from E. coli, B. stearothermophilus, Mtb and M. smeg.*** Amino acids colored in red are those deleted in the referenced studies. The conserved Tudor domain identified by Sanders, et al. is highlighted in blue.

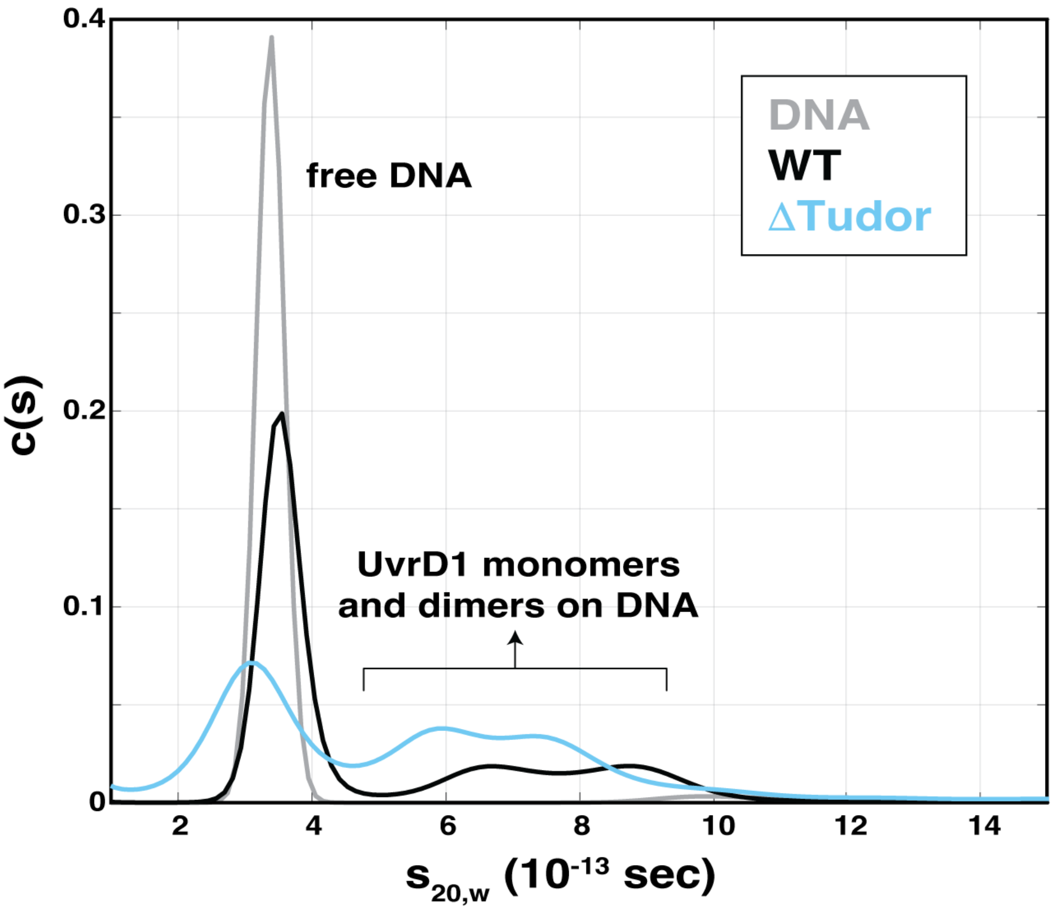


***Figure S8: Delta Tudor domain UvrD1 still binds DNA as a mixture of monomers and dimers in the absence of DTT.*** AUC velocity sedimentation data for labeled DNA alone (gray), WT UvrD1 (black), and ΔTudor UvrD1 (cyan). Only complexes containing the Cy5-labeled DNA were observed in this experiment.

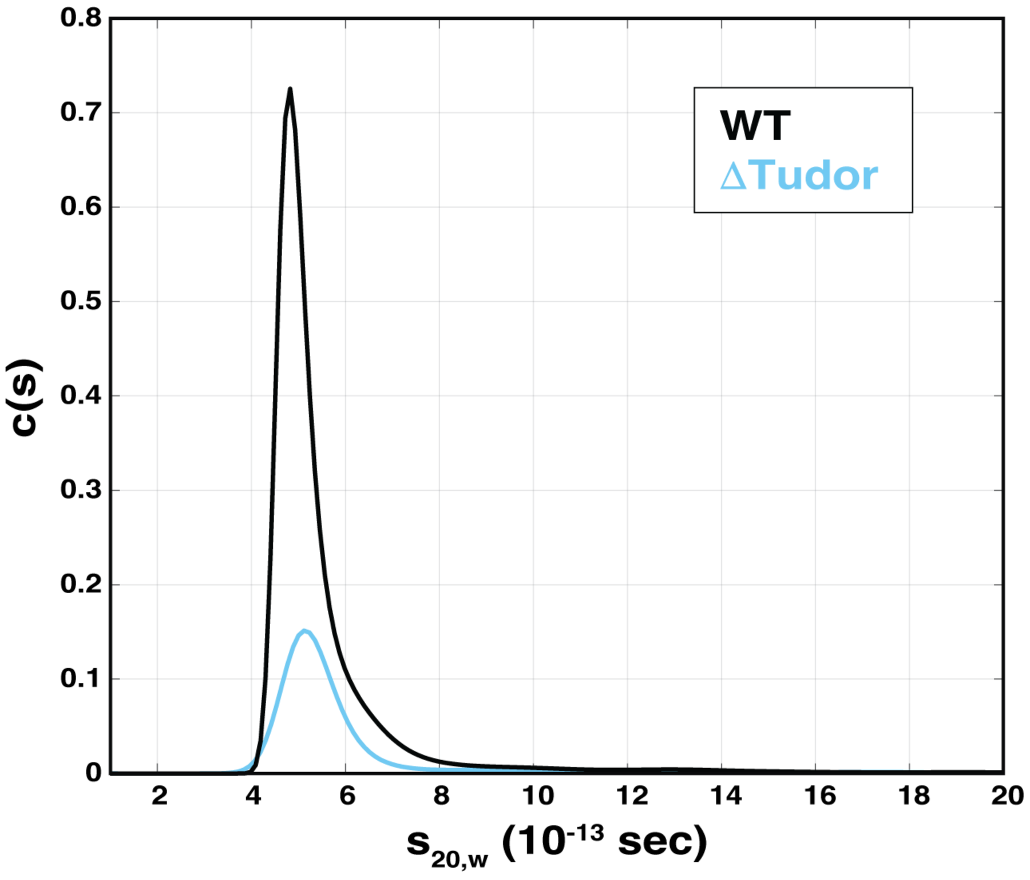


***Figure S9: ΔTudor UvrD1 is monomeric in the presence of DTT.*** AUC velocity data for WT (black) and ΔTudor UvrD1 in presence of DTT (cyan).

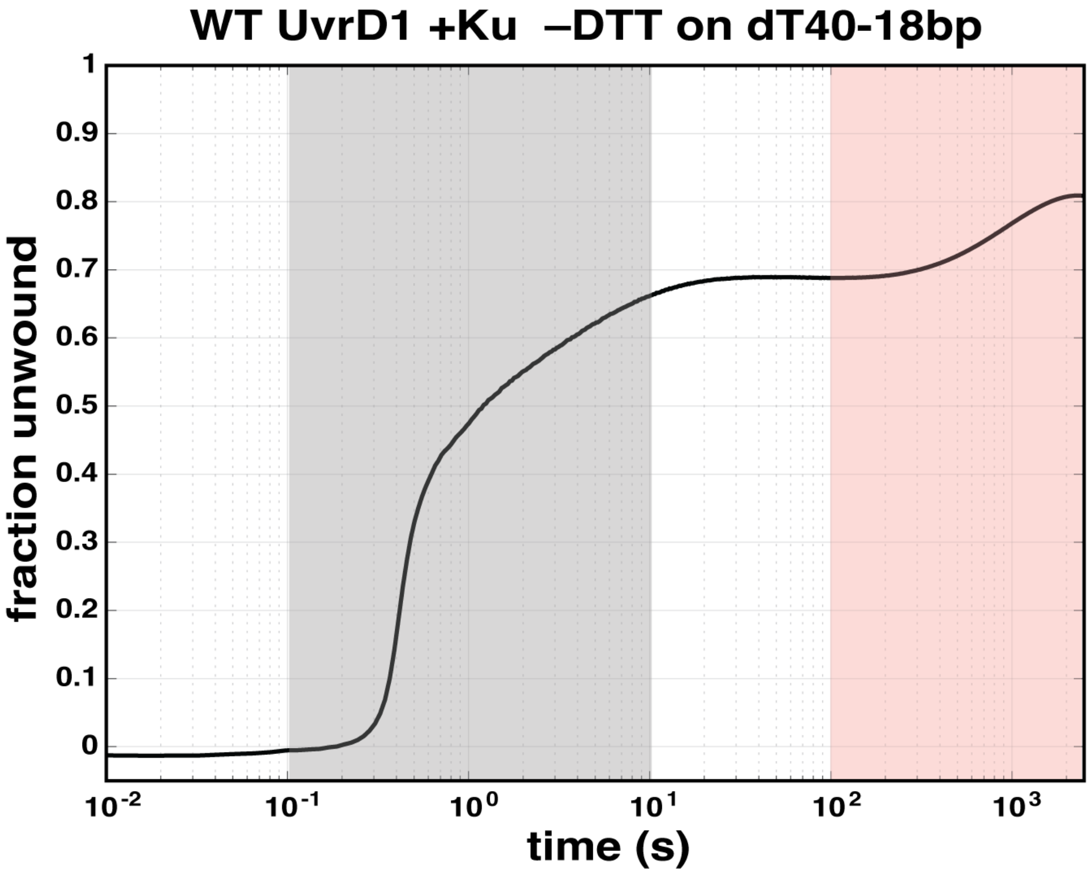


***Figure S10: Unwinding of WT UvrD1 with 18bp-dT40 in absence of DTT and presence of Ku on longer timescales:*** Multi-round DNA unwinding traces under oxidative conditions in the presence of Ku. The trace shows the sub-second unwinding of the UvrD1 dimer (gray shading) followed by a minutes-long second phase produced by monomeric UvrD1 in the presence of Ku (pink area).

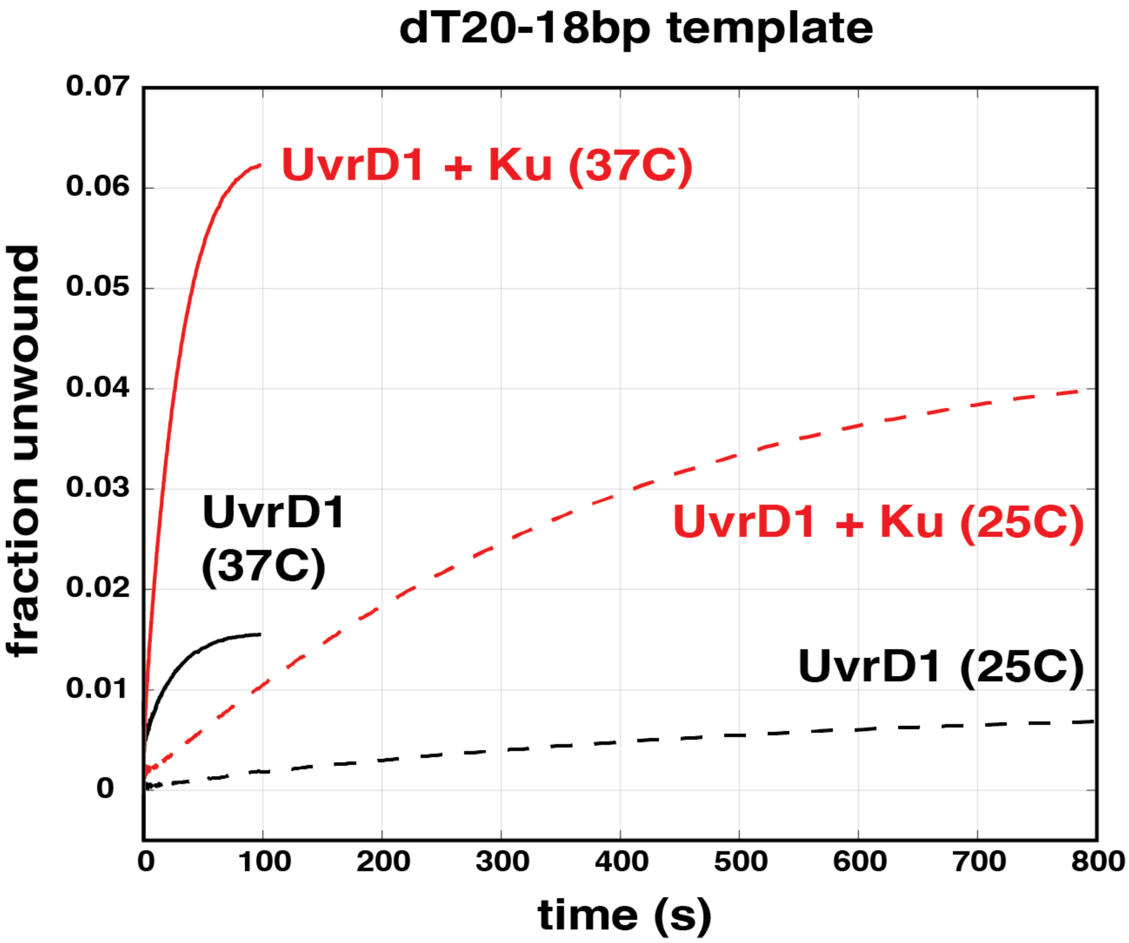


***Figure S11: The rate of unwinding of 2B UvrD1 is affected by temperature.*** Multi-round DNA unwinding traces of UvrD1 in the presence (red) and absence (black) of Ku at 37°C (solid) and 25°C (dashed). The rate of unwinding of an 18bp-dT20 substrate increases with temperature, but the fold-stimulation by Ku is comparable.

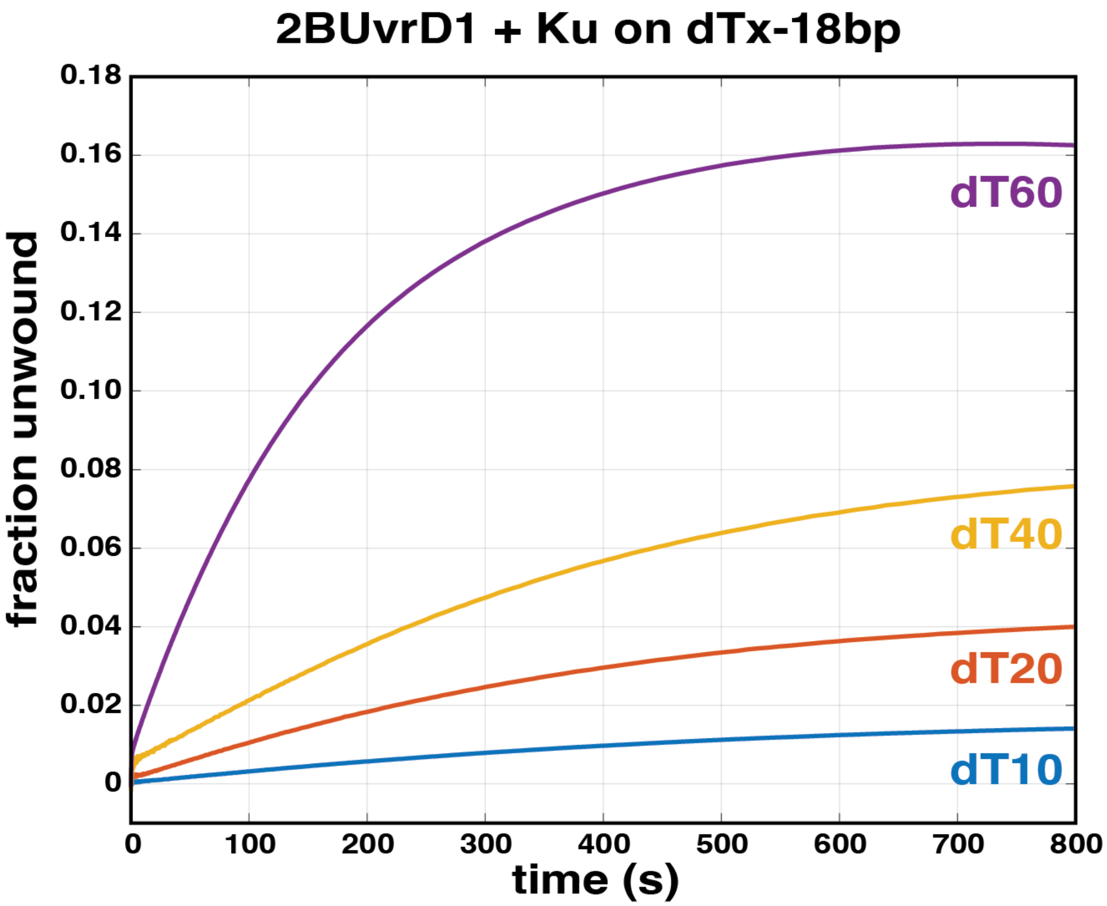


***Figure S12: Ku-dependent activation of monomer unwinding is single-strand tail dependent.*** Multi-round DNA unwinding traces in the presence of 2B UvrD1 on DNA substrates with increasing ssDNA tail lengths. The increase in fraction unwound with increasing tail length suggests that the ability to bind multiple monomers on a DNA substrate increases the efficiency of the unwinding reaction.

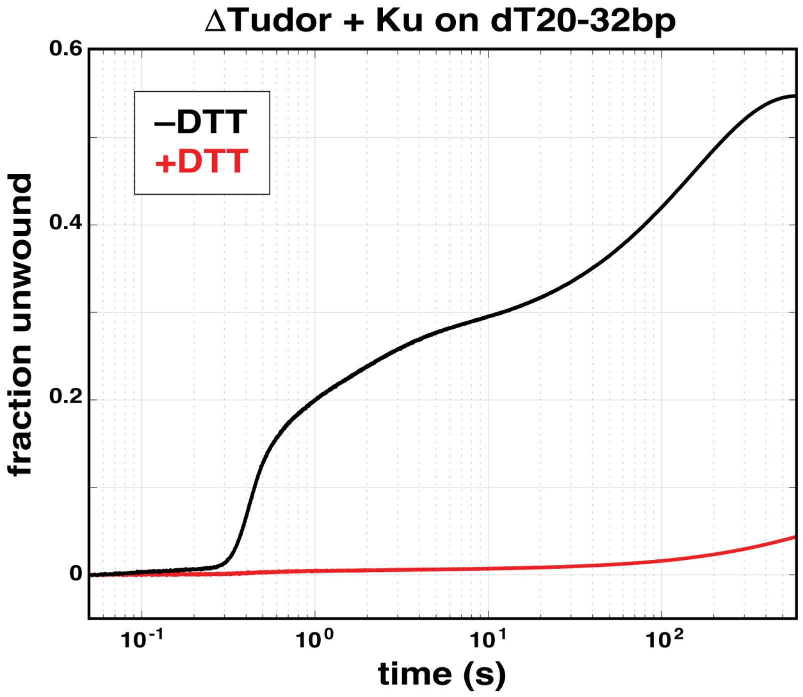


***Figure S13: Unwinding of ΔTudor UvrD1 on a 32bp-dT20 DNA substrate***. Multi-round DNA unwinding traces of ΔTudor UvrD1 in the presence (red) and absence (black) of DTT. The oxidation-dependent unwinding by UvrD1 dimers at short times can been observed in the black trace only while the slow phase due to the presence of monomers and stimulated by the removal of the Tudor domain can be observed in both traces.

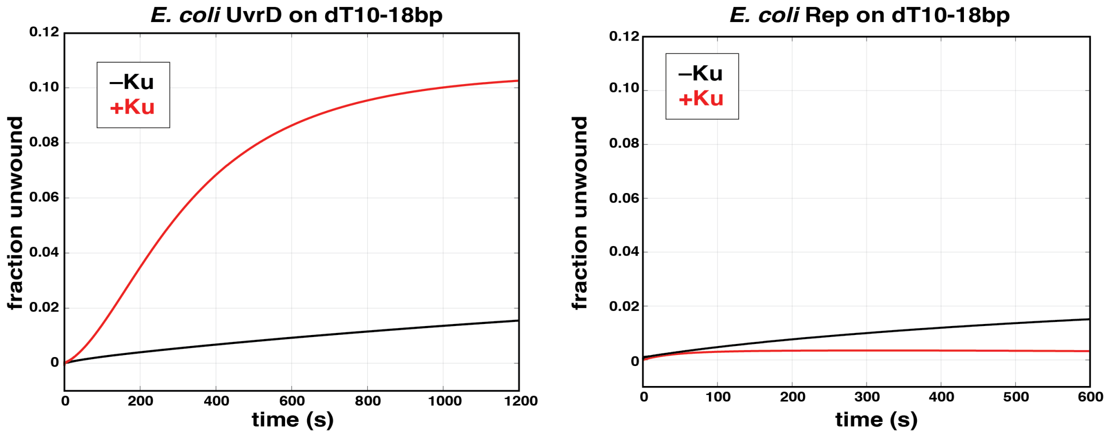


***Figure S14: Mtb Ku stimulates DNA unwinding by E. coli UvrD monomers but not Rep monomers.*** **(A)** Multi-round DNA unwinding traces of *E. coli* UvrD in the presence (red) and absence (black) of Ku. **(B)** Multi-round DNA unwinding traces of *E. coli* Rep in the presence (red) and absence (black) of Ku. *E. coli* UvrD contains a C-terminal Tudor domain whereas Rep does not further suggest that activation by Ku is through this Tudor domain.

*Supplemental Table 1: Sequences of single-stranded DNA used for unwinding and AUC*

| Lengt | Sequence | sequence type/assay |
|---|---|---|
| 18 | 5'GTT GGT CGG CAG CAG GGC-3' | ssDNA TRAP |
| 10bp- | 5'GCCTCGCTGCTTTTTGCAGCGAGGCTTTTTTTTTTTTTTTTTTTT TTTTTTTTTTTT 3' | hairpin TRAP |
| 52 | 5'-FAM GGA ATG TAA CCA TCG TTG GTC GGC AGC AGG GC dT(2 | Quenching assays |
| 32 | 5'- GCC CTG CTG CCG ACC AAC GAT GGT TAC ATT CC 3' | |
| 32 | 5'-FAM GGA ATG TAA CCA TCG TTG GTC GGC AGC AGG GC | |
| 20 | 5'-FAM dT(20)3' | |
| 38 | 5'-FAM GTT GGT CGG CAG CAG GGC dT(20)-3' | |
| 18 | 5'-GCC CTG CTG CCG ACC AAC -3' | |
| 39 | 5'-cy5 GTT GGT CGG CAG CAG GGC dT(40)-3' | Helicase assays |
| 52 | 5'-cy5 GTT GGT CGG CAG CAG GGC dT(20)-3' | |
| 32 | 5'-cy5 GGA ATG TAA CCA TCG TTG GTC GGC AGC AGG GC dT(20 | |
| | 5'-cy5 3' GCC CTG CTG CCG ACC AAC GAT GGT TAC ATT CC (BH | |
| 78 | 5'-cy5 GTT GGT CGG CAG CAG GGC dT(60)-3' | |
| 18 | 5'-GCC CTG CTG CCG ACC AAC (BHQ_2)-3' | |
| 50 | 5'-CCT GCA CTA GCA/iCy5/GCA GCG GGA ATG TAA CCA TCG TT GGC AGC AGG GC-3' | Analytical ultracentrifugation |
| 50 | 5'-GCC CTG CTG CCG ACC AAC GAT GGT TAC ATT CCC GCT GCT AGT GCA GG-3' | |
| 90 | 5'-dT(20) CCT GCA CTA GCA/iCy5/GCA GCG GGA ATG TAA CCA TTG GTC GGC AGC AGG GC dT(20)-3' | |
| 38 | 5'-cy5 GTT GGT CGG CAG CAG GGC dT(20)-3' | |
